# Activated fatty acid synthesis pathway in macrophages propagates pathogenic fibroblast expansion after myocardial infarction

**DOI:** 10.1101/2025.10.10.681697

**Authors:** Samreen Sadaf, Sathish Babu Vasamsetti, Imran Jamal, Ebin Johny, Khadeja-tul Kubra, Shagufta Haque, Araaf Mannan, Babak Razani, Srilakshmi Chaparala, Satoshi Okawa, Partha Dutta

## Abstract

Metabolic pathways, such as fatty acid oxidation and oxidative phosphorylation, can modulate inflammatory cells. However, little is known about the effects of the fatty acid synthesis pathway in macrophages on inflammation and cardiac remodeling after myocardial infarction (MI). Using spatial metabolomics, here we show that cardiac macrophages residing in the infarct synthesize *de novo* fatty acids and increase the production of fatty acid enzymes including ACLY and FASN. Mice deficient in myeloid *Acly* and *Fasn* have improved cardiac function after MI and reduced fibrosis. Combining Cleavage Under Targets and Release Using Nuclease (CUT&RUN), RNA sequencing analysis of *Acly^−/−^*macrophages, and macrophage-specific *in vivo* gene silencing, we demonstrate that ACLY acetylates the promoter region of the upstream regulator *Krt17*, which drives the production of pro-fibrotic cytokines, including IL-33. Single-cell RNA sequencing of cardiac fibroblasts shows that the expansion of a population of fibroblasts (Fibroblast 5) expressing high levels of extracellular matrix genes after MI is confined in the absence of macrophage *Acly*. Finally, the analysis of spatial multi-omics data of human hearts with MI uncovers myofibroblasts with the Fibroblast 5 gene signature. These myofibroblasts are located near cardiac macrophages expressing high levels of ACLY. In summary, we show that macrophage ACLY and FASN are deleterious in MI pathogenesis.

## Introduction

Cellular metabolic pathways support various biological functions including cell growth and survival. In recent years, several metabolic pathways have been involved in shaping immunity^1^. The contributions of glycolysis, fatty acid oxidation, and the Krebs cycle in the activation and modulation of immune cell functions, including leukocyte migration and recruitment, antigen presentation, and inflammatory cytokine production, are well established^1, 2, 3, 4, 5, 6, 7, 8^. Furthermore, inflammatory cell activity after myocardial infarction can be modulated by many metabolic pathways, including oxidative phosphorylation and fatty acid oxidation^9 10^. In the context of cardiovascular diseases, inflammatory cues and altered nutrient availability following ischemia contribute to metabolic shifts in immune cells that influence the course of disease pathology^11^. However, the significance of the fatty acid synthesis pathway, particularly the involvement of the enzymes in this pathway, in inflammation is not well understood.

MI leading to heart failure is a primary cause of morbidity and mortality in many developed countries. Although most MI patients survive the initial heart injury due to cutting-edge procedures like percutaneous coronary intervention, long-term morbidity and death from reinfarction and heart failure remain serious issues. Reinfarction and heart failure have been linked to excessive inflammation in MI patients^12,13,14^. Due to residual inflammation, a subset of individuals has significant cardiovascular mortality despite rigorous cholesterol-lowering therapy^15^. MI results in tissue injuries that entail immune cell influx to the infarcted region. This sets off an inflammatory reaction, followed by the repair of the wound and eventually scar formation.

Published studies demonstrated that cardiac macrophages play a central role in lingering inflammation and cardiac remodeling post MI^16,17,18 19^. Activation of immune cells^20^, such as macrophages and T cells, in diseases is concomitant with their epigenetic changes. Many enzymes involved in various metabolic pathways alter the epigenome via metabolic substrates. For instance, acetyl-CoA, a metabolite in the fatty acid synthesis pathway and produced by the cleavage of citrate by ATP citrate lyase (ACLY), is delivered to the nucleus where histone acetyltransferases use it as a substrate^21^. Additionally, ACLY is a known modulator of histone acetyltransferase activity^22^. Acetyl-CoA and histone acetyltransferase drive histone acetylation, which govern cellular processes by facilitating the access of transcription factors to histones. Congruently, histone acetyltransferase p300 induces IL-9 production and pulmonary inflammation^23^, and histone acetylase inhibition ameliorates left ventricular remodeling in a mouse model of diastolic dysfunction^24^. Thus, ACLY inhibition is a potential approach to modulate immune cell function by restricting histone acetylation. Surprisingly, whether ACLY governs cardiac macrophage-mediated inflammation and cardiac remodeling after myocardial infarction (MI) has not been explored. Moreover, post-MI epigenetic changes in these macrophages are not understood.

The goals of this study were: 1. To characterize *de novo* fatty acid synthesis in cardiac macrophages after MI, 2. To understand the significance of fatty acid synthesis enzymes expressed by cardiac macrophages in MI-induced cardiac remodeling, 3. To evaluate epigenetic alterations in cardiac macrophages post-MI, and 4. To examine the impact of macrophage fatty acid synthesis genes on infarct myofibroblast production. We demonstrated *de novo* synthesis of fatty acids and elevated production of the enzymes in the fatty acid synthesis pathway in cardiac macrophages after MI. Myeloid-specific *Acly* and *Fasn* deficient mice reduced fibrosis and improved cardiac function following MI. MI triggered the emergence of Fibroblast 5, a pathogenic fibroblast subset, in the heart following MI. Deletion of *Acly* in macrophages significantly depleted this fibroblast population. Mechanistically, ACLY promoted H3K27 acetylation of the promoter region of *Krt17*, which controlled pro-fibrotic cytokine production and cardiac fibrosis. Finally, patients with MI also harbored Fibroblast 5, which were located near ACLY^high^ cardiac macrophages in the heart.

## Results

### Myocardial infarction increases *de novo* fatty acid synthesis in cardiac macrophages

To characterize *de novo* fatty acid production in cardiac macrophages following MI, we performed spatial Matrix Assisted Laser Desorption/Ionization mass spectrometry imaging (MALDI-MSI) on sections obtained from hearts collected 28 days post-surgery from mice provided with deuterated water. After MALDI-MSI, we stained the tissue sections for CD68, a macrophage marker and superimposed the MALDI and immunofluorescence images to obtain the m/z ratios of fatty acids in macrophage clusters (**Figure 1A**). We found elevated levels of total fatty acids, such as palmitate, linolate, oleate, stearate, and arachidonate in macrophages present in the infarct area compared to the remote zone in the heart of mice with MI and the heart of sham-operated control mice (**Figure 1B-D and S1A**). We also observed increased deuterated fatty acids in the infarct, suggesting heightened fatty acid production in macrophages after MI. Total palmitate, oleate, stearate, and deuterated arachidonate were also increased in the remote zone of MI heart as compared to control hearts (**Figure 1B-D and S1A**). Next, we detected the enzymes required for fatty acid synthesis, including ATP citrate lyase (ACLY), acetyl-CoA carboxylase (ACC), and fatty acid synthase (FASN) by confocal microscopy. ACLY, ACC and FASN protein levels increased significantly in cardiac macrophages following MI in both mouse and human hearts (**Figure 1E and S1B**). Overall, these results suggest an increase in *de novo* fatty acid synthesis in cardiac macrophages after MI

**Figure 1:**
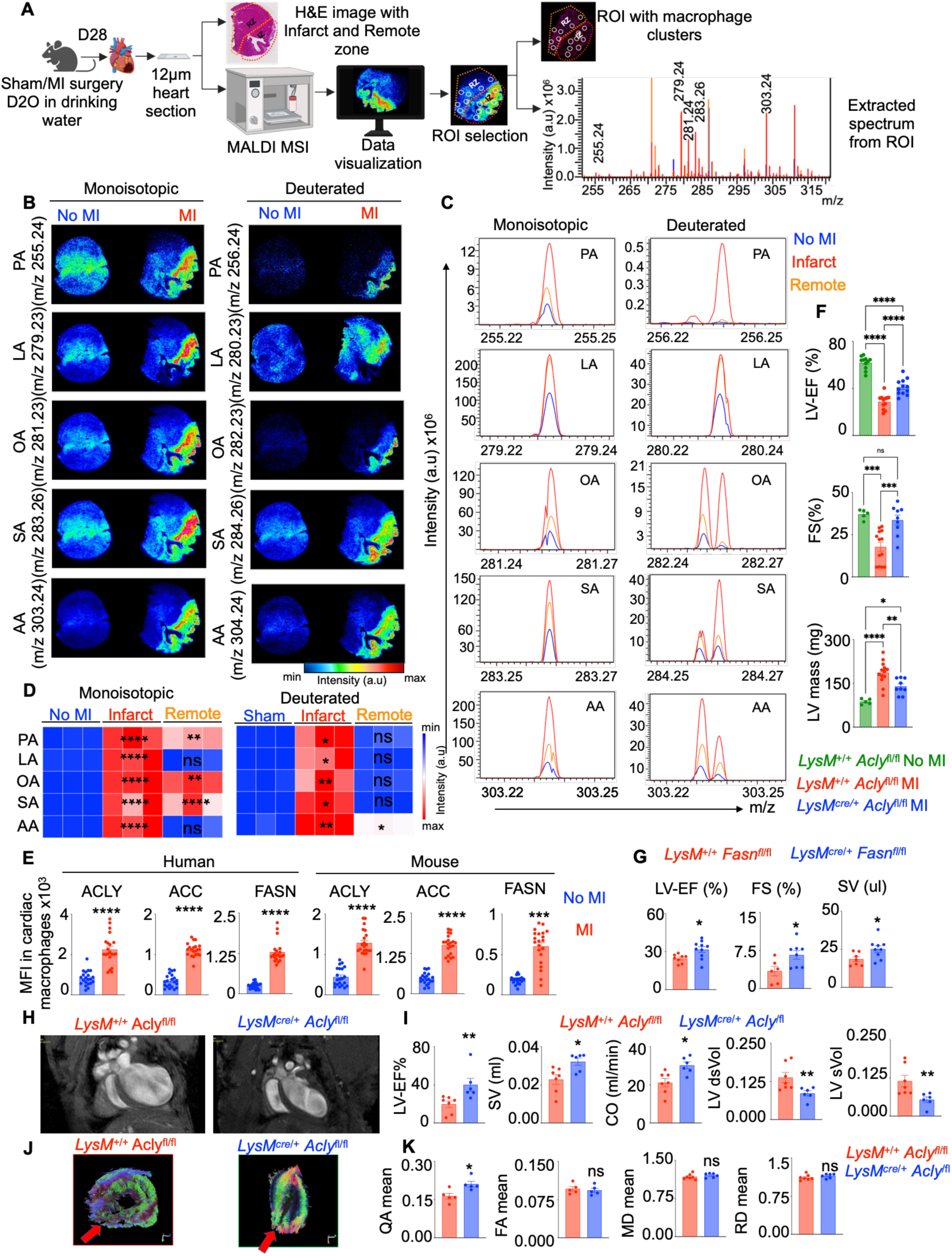
Fatty acid synthesis is augmented in cardiac macrophages after MI. **A.** The schematic diagram of the experimental design to quantify fatty acids in cardiac macrophages using MALDI is shown. **B-D.** The molecular images (**B**), representative single spectra (**C**), and heatmaps **(D)** of monoisotopic and D_2_O-incorporated palmitic acid (PA), linoleic acid (LA), oleic acid (OA), stearic acid (SA), and arachidonic acid (AA) acquired from macrophage clusters in the heart sections of C57BL/6 mice with sham and MI surgeries are provided. (n=3/group). **E.** Quantification of macrophage ATP citrate lyase (ACLY), acetyl-CoA carboxylase (ACC), and fatty acid synthase (FASN) mean fluorescence intensities (MFI) in control and MI patients and mice (day 28) are calculated by confocal microscopy (n=3-5/group). Each datapoint represents a tissue section. **F-G.** Left ventricular ejection fraction (LV-EF), left ventricle mass (LV mass), fractional shortening (FS), and stroke volume (SV) were calculated by echocardiography 28 days after MI in *LysM*^+/+^ *Acly*^fl/fl^ and *LysM^cre/+^ Acly*^fl/fl^ mice (n=5-13/group) (**F**) and in *LysM*^+/+^ *Fasn^fl^*^/fl^/ *LysM^cre/+^ Fasn*^fl/fl^ mice (n=6-10/group) (**G**). **H.** Representative four-chamber view by cine magnetic resonance images 28 days post MI is shown. **I.** Various cardiac parameters were assessed by cardiac magnetic resonance imaging (MRI) on 28 days after MI (n=5-7/group). **J-K.** High resolution diffusion tensor imaging (DTI) tractography in short axis view (**J**) and various DTI parameters (**K**) including quantitative anisotropy (QA), fractional anisotropy (FA), mean diffusivity (MD), and radial diffusivity (RD) on 28 days after MI are provided (n=5-7/group). The data are expressed as mean ± SEM and from 3–5 independent experiments. One way ANOVA for 1F and The Mann–Whitney U test for the other figures were used. \**p* < 0.05, \*\**p* < 0.01, \*\*\**p* < 0.001, \*\*\*\**p* < 0.0001.

### Myeloid-specific depletion of *Acly* and *Fasn* improves cardiac function following MI

To understand the impact of elevated levels of these enzymes after MI, we generated mice that lack *Acly* and *Fasn* in myeloid cells, namely *LysM^Cre/+^Acly^fl/fl^* and *LysM^Cre/+^Fasn^fl/fl^*, respectively (**Figure S1C-D**). *LysM^Cre/+^Acly^fl/fl^* mice had higher ejection fraction and fractional shortening and lower LV mass, diastolic volume, and systolic volume, indicating their improved cardiac function following MI (**Figure 1F and S1E-F**). However, we did not find any change in the echocardiographic parameters like ejection fraction, stroke volume, fractional shortening, and diastolic and systolic volume in mice without MI (**Figure S1G)**. Similar to *LysM^Cre/+^Acly^fl/fl^* mice with MI, *LysM^Cre/+^Fasn^fl/fl^*mice had higher ejection fraction, fractional shortening, and stroke volume 28 days following MI compared to the littermate control mice (**Figure 1G and S1H**). Assessment of the hearts of *LysM^Cre/+^Acly^fl/fl^* mice after MI by magnetic resonance imaging (MRI) confirmed improved cardiac function as evidenced by enhanced ejection fraction, stroke volume, and cardiac output (**Figure 1H-I** and **S1I**). Myeloid deficiency of *Acly* also diminished left ventricular diastolic and systolic volumes, indicating curtailed ventricular dilation following MI. Finally, diffusion tensor imaging of the explanted hearts from these mice revealed improved structure, organization, and integrity of the heart in the absence of myeloid *Acly* (**Figure 1J-K**). Since inflammatory cells determine post-MI cardiac remodeling^25^, we enumerated leukocyte populations, including macrophages, monocytes, neutrophils, B cells, T cells, and dendritic cells, in the heart, blood, bone marrow, and spleen of *LysM^Cre/+^Acly^fl/fl^*, *LysM^Cre/+^Fasn^fl/fl^*, and their littermate control mice. Surprisingly, we did not see any significant change in immune cell numbers in these mice (**Figure S2A and S3**). Interestingly, we saw a significant downregulation in inflammatory gene expression in *Acly^−/−^* cardiac macrophages isolated after MI, implying that ACLY does not affect the number of macrophages, but escalates inflammation mediated by cardiac macrophages (**Figure S2B**).

### ACLY elevates pro-fibrotic genes in cardiac macrophages after MI

To identify the genes regulated by ACLY in cardiac macrophages residing in the infarct, we performed bulk RNA sequencing in these cells isolated from *LysM^+/+^Acly^fl/fl^* and *LysM^Cre/+^Acly^fl/fl^* mice on day 28 after MI. The PCA plot (**Figure 2A**) and heatmap (**Figure 2B**) show that the *Acly* deficiency significantly altered the gene expression prolife in cardiac macrophages after MI. Several genes implicated in fibrosis and fibrosis modulation^26, 27^ were significantly altered in the absence of *Acly*, such as *Il10, Il33*, *Adamts5, Flt1, Ctsk, Cidec, Cav1, Slc6A4*, and *Csrp3* (**Figures 2B and 2C**). In agreement with this observation, Ingenuity Pathway Analysis (IPA) revealed several fibrosis pathways, such as “fibrosis of heart” and “quantity of connective tissue”, to be downregulated in *Acly*-deficient macrophages (**Figure 2D).** Interestingly, *Acly* deficiency enriched the genes driving the pathways “cardiac muscle function” and “cardiac contractility” (**Figure S4A**). Finally, IPA analysis uncovered ten upstream regulators predicted to drive the transcriptomic differences in *Acly*^−/−^ cardiac macrophages compared to their *Acly*^+/+^ counterpart (**Figure S4B**). Among these upstream regulators, *Egr1*, *Ikbkb*, *Nlrp3*, *Krt17*, *Ikbkg*, and *Chuk* were significantly inhibited in *Acly*-deficient cardiac macrophages (**Figures 2E-F and S4B**). Altogether, these results suggest that following MI, ACLY raises pro-fibrotic genes in cardiac macrophages.

**Figure 2:**
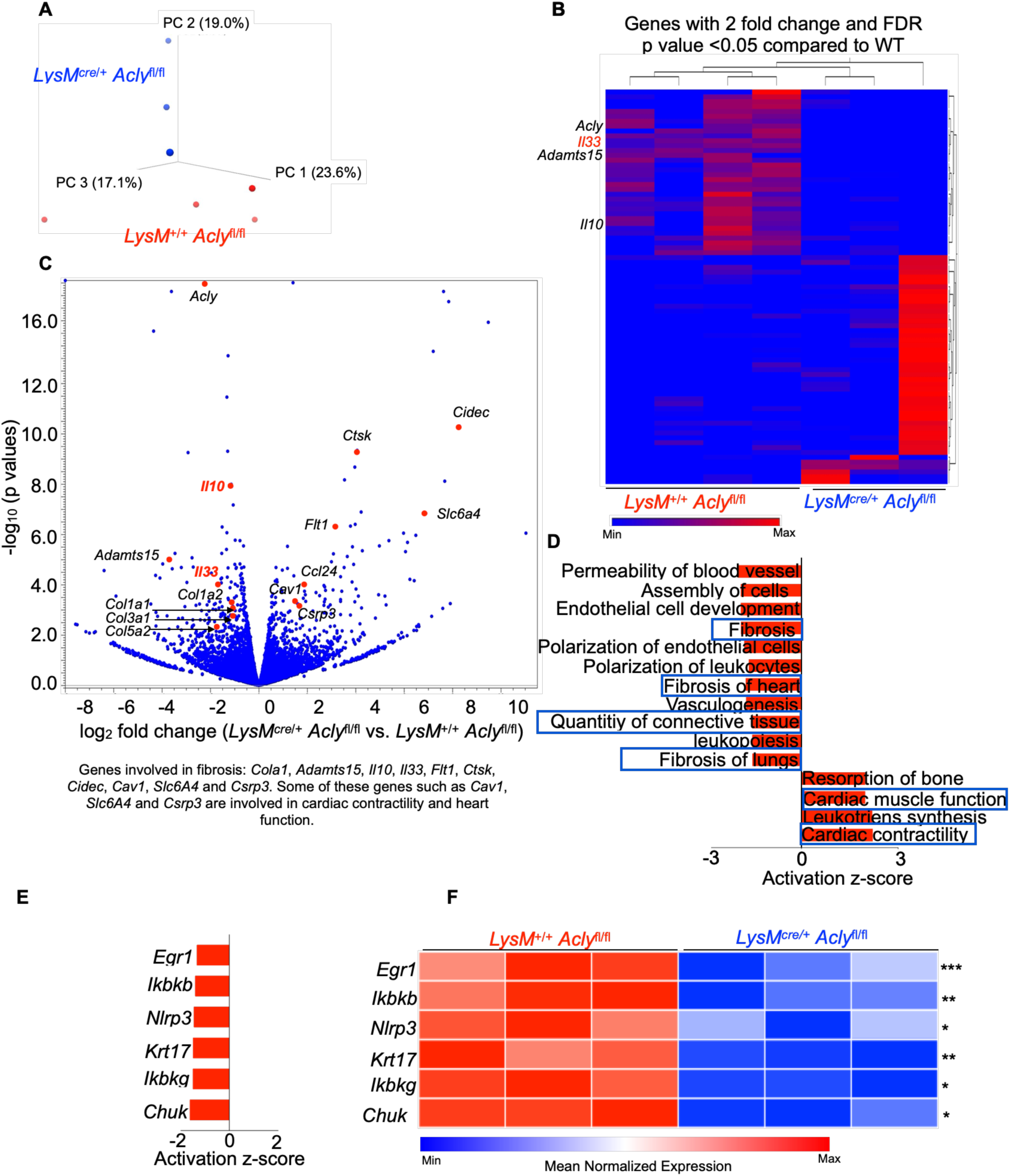
ACLY elevates pro-fibrotic genes in cardiac macrophages after MI. Cardiac macrophages were isolated from *LysM*^+/+^ *Acly*^fl/fl^ and *LysM^cre/+^ Acly*^fl/fl^ mice on day 28 after MI, and bulk RNA sequencing was performed. **A-C** Principal component analysis (PCA) **(A),** heatmap (**B**), and volcano plot (**C**) show the transcriptional differences in *LysM^cre/+^ Acly*^fl/fl^ vs *LysM*^+/+^ *Acly*^fl/fl^ cardiac macrophages (n=3). **D.** An Ingenuity Pathway Analysis of the differentially expressed genes (n=3) was carried out. **E**. The upstream regulators downregulated in *Acly*-deficient cardiac macrophages are identified (n=3). **F.** The heatmap depicts the expression of the upstream regulator in cardiac macrophages by qPCR on 28 days after MI surgery (n=10-12/group). qPCR data are expressed as mean ± SEM from 3–5 independent experiments. The Mann-Whitney U test was performed in the Fig 2F. \**p* < 0.05, \*\**p* < 0.01, \*\*\**p* < 0.001.

### Macrophage ACLY exacerbates cardiac fibrosis after MI

MI often leads to cardiac remodeling and heart failure due to uncontrolled myofibroblast proliferation and fibrosis^28^. As we observed an enrichment of fibrosis pathways in the presence of *Acly* in cardiac macrophages after MI, we next explored the effect of macrophage ACLY on cardiac fibrosis. Magnetic resonance imaging (MRI) revealed a significant diminution in left ventricular extracellular volume (LV-ECV), which indicates interstitial myocardial fibrosis in *LysM^Cre/+^Acly^fl/fl^*as compared to the control *LysM^+/+^Acly^fl/f^* (**Figure 3A**). Consistently, we observed a significant reduction in myocardial fibrosis quantified by Masson’s trichome staining in the absence of myeloid *Acly* (**Figure 3B**). Using qPCR, we also observed a significant reduction in the expression of fibrosis genes like *Col1a1*, *Col1a2*, *Sparc*, and *Fibronectin* in the infarcted area of *LysM^Cre/+^Acly^fl/fl^* mice (**Figure 3C**). The levels of matrix metalloproteinases involved in ventricular dilation^29^ were decreased. In line with these findings, fibroblasts sorted from these mice^30, 31, 32^ (**Figure S5A**) had reduced collagen synthesis genes and the cell proliferation gene mKi67 (**Figure 3D**). Concomitant with suppressed IL-33 production by *Acly*^−/−^ cardiac macrophages (**Figure 2B**), we detected a significant reduction in the gene expression of IL33 receptor (*Il1rl1*), also known as ST2, in fibroblasts sorted from *LysM^Cre/+^Acly^fl/fl^*mice (**Figure 3D**). We next quantified fibroblast proliferation using vimentin as a resting fibroblast marker, α-smooth muscle actin (α-SMA) as a marker for myofibroblasts, and Ki67 as a proliferation marker. Mice lacking myeloid *Acly* had reduced numbers of myofibroblasts and proliferating fibroblasts (**Figure 3E**). To further confirm the impact of macrophage ACLY on fibroblasts, we cocultured bone marrow-derived macrophages (BMDM) from *LysM^+/+^Acly^fl/f^*and *LysM^Cre/+^Acly^fl/fl^* mice with cardiac fibroblasts isolated from C57BL/6 mice (**Figure S5B-D**). Fibroblasts co-cultured with *Acly* KO BMDM were less proliferative (**Figures 3F-G and S5D**). The sorted cardiac fibroblasts from *Acly* KO BMDM co-cultures also had a significantly reduced expression of the collagen synthesis genes and the IL33 receptor (**Figure 3H**). Similar to myeloid *Acly*-deficient mice, *LysM*^cre/+^ *Fasn*^fl/fl^ mice exhibited dampened myofibroblast expansion and proliferation in the heart after MI (**Figure S5E**). Together, these results imply a critical role of the macrophage fatty acid synthesis enzymes ACLY and FASN in post-MI cardiac fibrosis.

**Figure 3:**
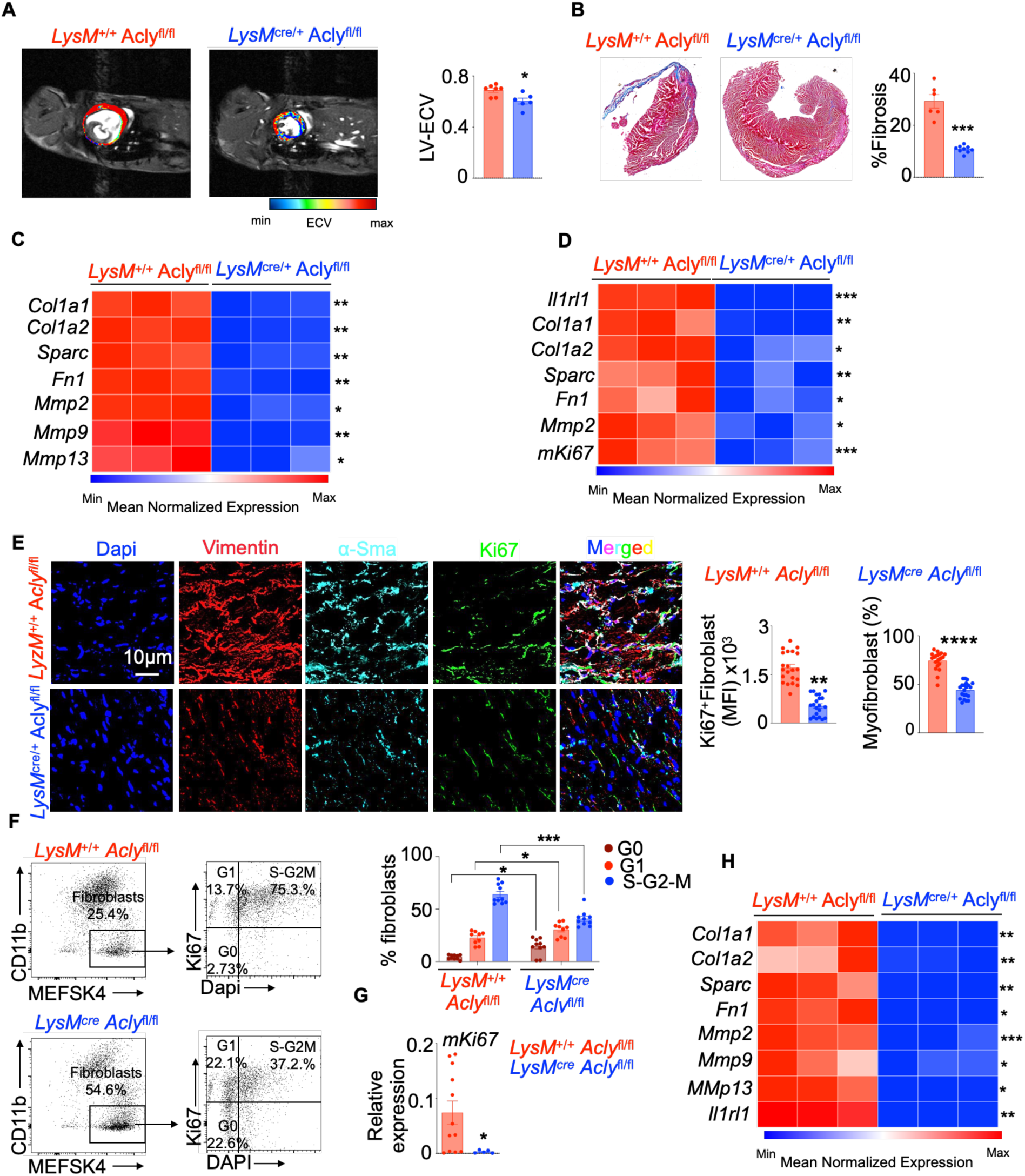
Macrophage ACLY exacerbates cardiac fibrosis after MI. **A.** MRI was employed to generate extracellular volume (ECV) maps by imaging enhanced gadolinium (Gd) accumulation in extracellular regions in the hearts of *LysM*^+/+^ *Acly*^fl/fl^ and *LysM^cre/+^ Acly*^fl/fl^ 28 days after MI (n=5/group). **B.** Masson trichrome staining and its quantification are shown in the heart on 28 days after MI (n=6-9/group). **C-D.** The heatmap depicts the expression of the genes involved in collagen synthesis, processing, and degradation in the heart (**C**) and sorted cardiac fibroblasts (**D**) on day 28 after MI (n=5-7/group). **E.** Representative confocal microscopy images for vimentin, α-SMA, and Ki-67 in the heart, and the quantification of Ki-67^+^ fibroblasts and frequency of myofibroblasts are included. (n=5-7/group). **F.** C57BL/6 cardiac fibroblasts were cocultured with *LysM*^+/+^ *Acly*^fl/fl^ or *LysM^cre/+^ Acly*^fl/fl^ BMDM, and their cell cycle status was evaluated by flow cytometry (n=5-12/group). **G-H.** qPCR quantification of *mKi67* (**G**) and the heatmap of the fibrosis pathway genes and *Il1rl1* (**H**) in C57BL/6 cardiac fibroblasts cocultured with *LysM^+/+^ Acly*^fl/fl^ or *LysM^cre/+^ Acly*^fl/fl^ BMDM (n=5-12/group). The data are expressed as mean ± SEM and pooled from 3–5 independent experiments. One way ANOVA was performed for Fig 3F, and The Mann-Whitney U test was used for the other data\**p* < 0.05, \*\**p* < 0.01, \*\*\**p* < 0.001, \*\*\*\**p* < 0.0001.

### Krt17 regulated by macrophage Acly precipitates fibrosis

To delineate the mechanisms of ACLY-mediated cardiac fibrosis, we silenced the upstream regulators *Egr1, Ikbkb, Nlrp3, Krt17, Ikbkg,* and *Chuk* in BMDM (**Figure S6A**), which we found to be controlled by ACLY (**Figures 2E-F and S4B**). We examined the expression of profibrotic and proreparative genes including *Il13, Il15, Chil3, Tgfb, Il4* and *Il15*. The silencing of all genes but *Chuk* reduced expression of the most of the profibrotic and proreparative genes (**Figure 4A**). Because ACLY links cell metabolism to histone acetylation, we measured H3K27ac in the promoter regions of these upstream regulators in the presence or absence of *Acly*. *Acly* deficiency curtailed H3K27ac in the *Nlrp3* and *Krt17* promoters but not in *Egr1, Ikbkg,* and *Ikbkb* promoters in macrophages (**Figure 4B**). An IPA analysis revealed that Krt17 controlled many profibrotic cytokine and chemokine genes including *Ccl17, Ccl22, Ccl24, Cxcl2*, and *Il10* (**Figure S6B**). To understand if Krt17 orchestrates fibrosis, we silenced Krt17 *in vivo* in tissue macrophages in C57BL/6 using lipidoid nanoparticles as we did previously^33^. We saw a significant improvement in cardiac function such as left ventricular ejection fraction, systolic volume, and cardiac output after si*Krt17* treatment (**Figure 4C**), partially phenocopying the effects of *Acly* depletion. Immunophenotyping showed that the *siKrt17-*injected mice harbored significantly fewer macrophages and T cells in the heart and more in the spleen compared to the control mice (**Figure S6C**), suggesting a reduced recruitment of immune cells to the heart post-injury. Cardiac myofibroblast proliferation and frequency were suppressed in the infarct of *sikrt17*-treated mice compared to the mice injected with *siControl* (**Figure S6D)**. In line with these findings, cardiac macrophages isolated from the mice after *Krt17* silencing expressed reduced amount of the pro-fibrotic factors (**Figure 4D**), such as the pro-fibrotic cytokine *Il33,* which is also regulated by ACLY (**Figure 2B-C**). A similar downregulation of profibrotic genes was observed upon *in vitro* silencing of Krt17 in BMDM (**Figures 4E, S6E**). As we observed suppressed IL-33 production by *Acly*^−/−^ cardiac macrophages (**Figure 2B**) and a significant reduction in the gene expression of IL33 receptor (*Il1rl1*) by *Acly*^−/−^ fibroblasts (**Figure 3D**), we investigated if *Il33* is regulated by the upstream regulators. Indeed, we observed the silencing of *Egr1*, *Ikbkb*, *Krt17*, *Ikbkg*, and *Nlrp3* diminished Il33 expression by macrophages (**Figures 4E**). Moreover, *Acly*-deficient BMDM exhibited suppressed acetylation of the *Il33* promoter as estimated by ChIP-qPCR (**Figure 4F**). In line with the studies reporting pro-fibrotic effect of IL-33, the supplementation of this cytokine in *LysM^Cre/+^Acly^fl/fl^* mice restrained cardiac ejection fraction (**Figure 4G and S7A**) and enhanced cardiac fibrosis (**Figure 4H**) and fibroblast proliferation (**Figure S7B**) in these mice. In summary, these data indicate that Krt17 promotes cardiac fibrosis after MI.

**Figure 4:**
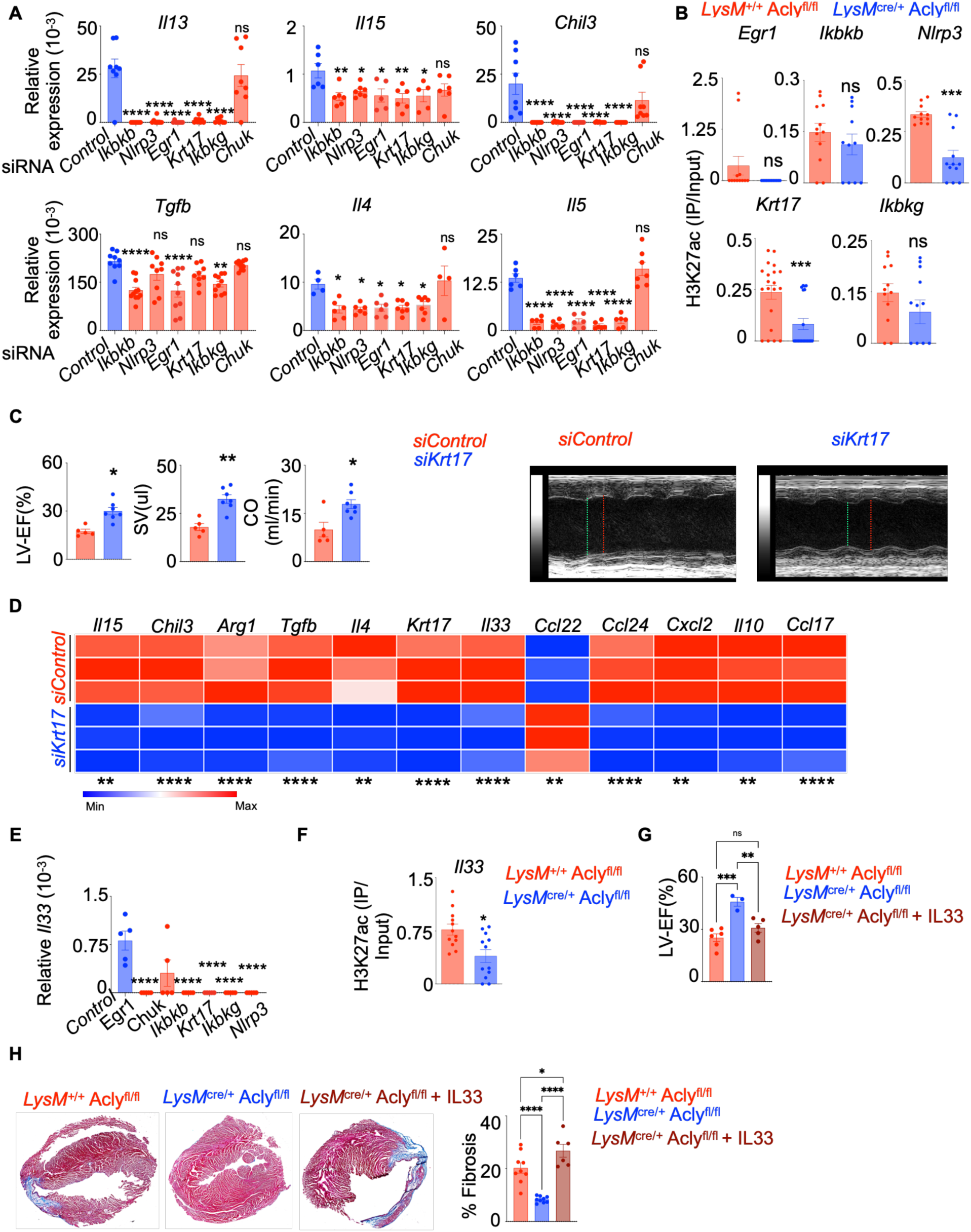
Krt17 regulated by macrophage ACLY precipitates fibrosis. **A.** The profibrotic genes (*Il13*, *Il15*, *Chil3*, *Tgfb*, *Il4*, and *Il5*) in BMDM treated with si*Control*, si*Ikbkb*, si*Nlrp3*, si*Egr1*, si*Krt17*, and si*Ikbkg* were quantified by qPCR. (n=6-10/group). **B.** ChIP-qPCR was conducted to discern the H3K27Ac mark in the promotor regions of *Egr1*, *Ikbkb*, *Nlrp3*, *Krt17,* and *Ikbkg* in *Acly*^+/+^ and *Acly*^−/−^ BMDM. (n=10-12/group). **C.** Left ventricular ejection fraction (LV-EF), stroke volume (SV), and cardiac output (CO) were measured by echocardiography in C57BL/6 mice after si*Control* and si*Krt17* treatment for 28 days post MI. (n=5-7/group). The images depict representative echocardiograms of the LV in M-mode in SAX view in mice treated with *siControl* and *siKrt17* till day 28 after MI (n=5-7/group). The green and red lines indicate LVID;s = left ventricular internal diameter in systole and LVID;d = left ventricular internal diameter in diastole respectively **D.** The heatmap displays the expression of the profibrotic genes in cardiac macrophages sorted from si*Control* and si*Krt17*-treated mice 28 days after MI. (n=5-7)/group. **E.** *IL33* expression in BMDM after silencing *Egr1*, *Chuk*, *Ikbkb*, *Krt17*, *Nlrp3*, and *Ikbkg* was ascertained by qPCR (n=4-5/group). **F.** The H3K27Ac mark in the promotor region of *Il33* in *Acly*^+/+^ and *Acly*^−/−^ BMDM was estimated by ChIP-qPCR. (n=10-12/group). **G-H.** LV-EF was evaluated by echocardiography (**G**) and Masson’s trichrome staining was performed to quantify cardiac fibrosis (**H**) in *LysM*^+/+^ *Acly*^fl/fl^ and *LysM^cre/+^ Acly*^fl/fl^ mice treatred with vehicle or IL-33 for 28 days following MI (n=9-6/group). The data are expressed as mean ± SEM and pooled from 3–5 independent experiments. One way ANOVA was performed for Fig 4A, 4E, 4G, and 4H, and The Mann-Whitney U test was used for the analysis of the rest of the data. \**p* < 0.05, \*\**p* < 0.01, \*\*\**p* < 0.001, \*\*\*\**p* < 0.0001.

### Macrophage ACLY imprints H3K27ac in the promoter regions of profibrotic genes

ACLY mediates histone acetylation. To discern if ACLY imprints unique epigenetic marks in macrophages, we performed CUT&RUN for H3K27ac, a gene activation mark, in *Acly*^+/+^ and *Acly*^−/−^ BMDM. We observed the H3K27 acetylation mark at different regions of genes (**Figure 5A**). This epigenetic mark was significantly restricted at promoter regions in *Acly*^−/−^ BMDM (**Figure 5B-C**). *Acly*^−/−^ macrophages had reduced H3K27ac mark in the promoters of *Ikbkb, Nlrp3,* and *Krt1*7 (**Figure 5D**), which are upstream regulators controlled by ACLY (**Figure 2E**), implying that the activation of these genes is instigated by ACLY-mediated acetylation. The promoter of *Il33*, a profibrotic cytokine downregulated in *Acly*^−/−^ cardiac macrophages (**Figure 2B-C**), also has diminished H3K27ac mark (**Figure 5E**). We next compared the genes having reduced H3K27ac as determined by CUT&RUN and the genes with suppressed expression assessed by bulk RNA seq in *Acly*^−/−^ cardiac macrophages. We found 18 common genes, such as *Ccr1, Sparc, Ccr5,* and *Fbxwr7,* which have been reported to be critical in inflammation and fibrosis^34 35 36^ (**Figure 5F and Table S1**). Pathway analysis of the common genes (**Figure 5G**) and the genes with reduced H3K27ac mark in the absence of *Acly* (**Figure 5H**) revealed several pathways involved in inflammation such as C-C Chemokine Receptor Activity, C-C Chemokine Binding, Transcription Regulator Activity, Double Stranded DNA Binding, and Regulation of Primary Metabolic Process. Finally, Motif Enrichment Analysis uncovered several transcription factors, such as KLF16, RREB1, and MZF1 (**Figure 5I**), which control fibrosis^37 38 39^. Altogether, these results convey that macrophage ACLY imprints the H3K27ac mark in the promoter regions of the profibrotic regulators.

**Figure 5:**
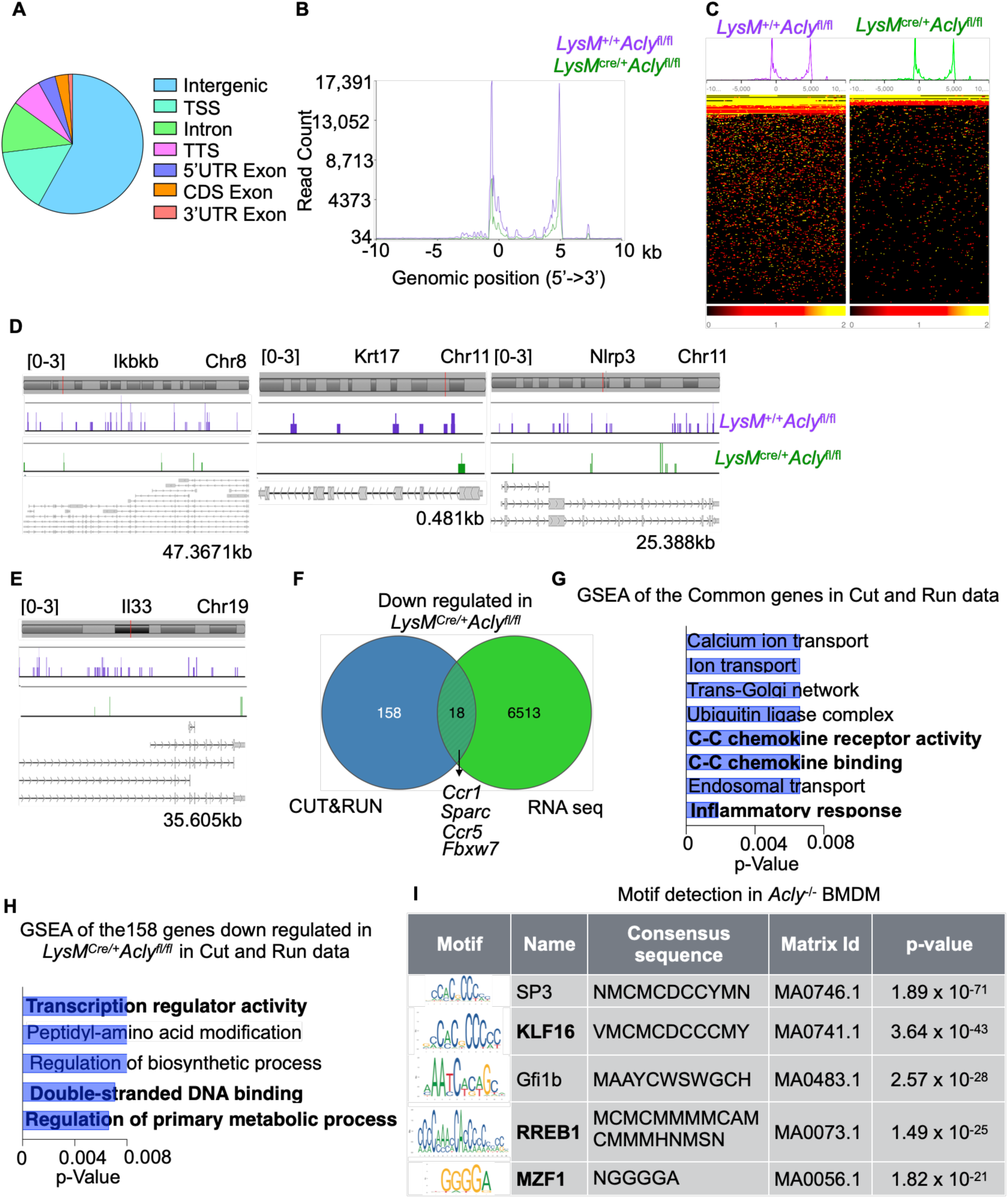
Macrophage ACLY imprints H3K27ac in the promoter regions of profibrotic genes. CUT&RUN was performed to detect genome wide H3K27Ac mark in BMDM obtained from *LysM*^+/+^ *Acly*^fl/fl^ and *LysM*^cre/+^ *Acly*^fl/fl^ mice. n=3/ group. **A**. H3K27ac distribution in the various parts of the genome is depicted. **B-C.** The intensity plot (**B**) and the heatmap (**C**) obtained from the CUT&RUN analysis display H3K27Ac enrichment around +/−10 kb of the transcription start sites. **D-E**. H3K27Ac peak distributions in the *Ikbkb, Krt17,* and *Nlrp3* genes (**D**) and the *Il33* gene **(E)** are shown. **F.** The 18 common genes are identified in 1. the genes having reduced H3K27ac mark in the CUT&RUN analysis in *LysM*^cre/+^ *Acly*^fl/fl^ BMDM and 2. the downregulated genes in *LysM*^cre/+^ *Acly*^fl/fl^ cardiac macrophages assessed by bulk RNA sequencing. **G-H.** Gene Set Enrichment analyses of the common genes obtained in F (**G**) and the genes exhibiting reduced H3K27Ac mark in *LysM*^cre/+^ *Acly*^fl/fl^ BMDM (**H**) were performed. **I.** The motifs were detected.

### Macrophage ACLY facilitates the expansion of Fibroblast 5 after MI

To elucidate the molecular mechanisms underlying the ACLY-mediated fibrosis in response to MI, we performed single cell RNA seq in cardiac fibroblasts isolated from *LysM^+/+^Acly^fl/fl^*and *LysM^Cre/+^Acly^fl/fl^* on day 28 after sham or MI surgeries. Our analysis revealed seven cardiac fibroblast subsets (**Figure 6A-B and Table S2**). Further characterization of these populations uncovered enrichment of Fibroblasts 3, 4, and 5 in the infarct after MI in the presence of myeloid *Acly* while fibroblast 1 and 2 predominated in the hearts of sham-operated mice. Interestingly, the depletion of myeloid *Acly* led to the shrinkage of Fibroblasts 3, 4, and 5 after MI (**Figures 6C-D and S8A-B**). Among these fibroblasts, Fibroblast 5 highly expressed the genes enriched in activated fibroblasts, myofibroblasts, and cardiac fibroblast secretome^31, 40, 41, 42 43 44 45^ (**Figure 6E-F, S8C-E and Table S3**). Pseudotime trajectory analysis of the fibroblast subsets revealed that Fibroblasts 1-2 are relatively immature fibroblasts while Fibroblasts 3-7 are mature fibroblasts (**Figure 6G-H**). Trajectory analysis of the genes in activated fibroblast, myofibroblasts, and cardiac fibroblast secretome further confirmed that these genes are the hallmark of Fibroblast 5 (**Figure S8F**). To find the genes regulated by myeloid ACLY in Fibroblast 5, we identified the common differentially expressed genes in these two comparisons: A) the upregulated genes in Fibroblast 5 in the infarct of *Acly*^+/+^ mice compared to Fibroblasts 1 and 2 in *Acly*^+/+^ sham-operated mice and B) the downregulated genes in Fibroblasts 1 and 2 in the infarct of *Acly*^−/−^ mice compared to Fibroblast 5 in the infarct of *Acly*^+/+^ mice (**Figure 6I**). This analysis revealed 204 common genes, which were, as expected, enriched in Fibroblast 5 (**Figure S8G and Table S4**). Furthermore, we discerned differentially expressed genes in Fibroblast 5 vs Fibroblasts 1 and 2 and observed several genes, such as *Comp*, *Ecrg4, Cthrc1, Ddah1, Fmod,* and *Cilp2*, to be upregulated (**Figure 6J, S8H and Table 5**). Pathway analysis with the differentially expressed genes revealed activation of actin cytoskeleton, wound healing, tumor microenvironment, Gp6 signaling, and fibrosis (**Figure 6K**). The pathway analysis also unveiled a set of activated transcription factors (TFs) regulating the expression of Fibroblast 5 genes (**Figure 6L**). qPCR on Cardiac fibroblasts isolated from the infarct area of *LysM^+/+^Acly^fl/fl^* mice validated the expression of these genes at high levels compared to the ones isolated from *LysM^Cre/+^Acly^fl/fl^* mice (**Figure 6M**). To further examine the contribution of macrophage ACLY in the upregulation of these transcription factors, we assessed their expression in cardiac fibroblasts cultured in the presence of *Acly*^+/+^ and *Acly*^−/−^ BMDM. The sorted cardiac fibroblasts cocultured in the presence of *Acly*^−/−^ BMDM displayed reduced expression of *Msx2, Tead2, Ccnd1, Mycn, Glis1, Snai1, and Tead4* (**Figure 6N**). Overall, these results suggest that the emergence of pathogenic Fibroblast 5, which is enriched in the genes expressed by myofibroblasts, after MI is myeloid ACLY-dependent.

**Figure 6:**
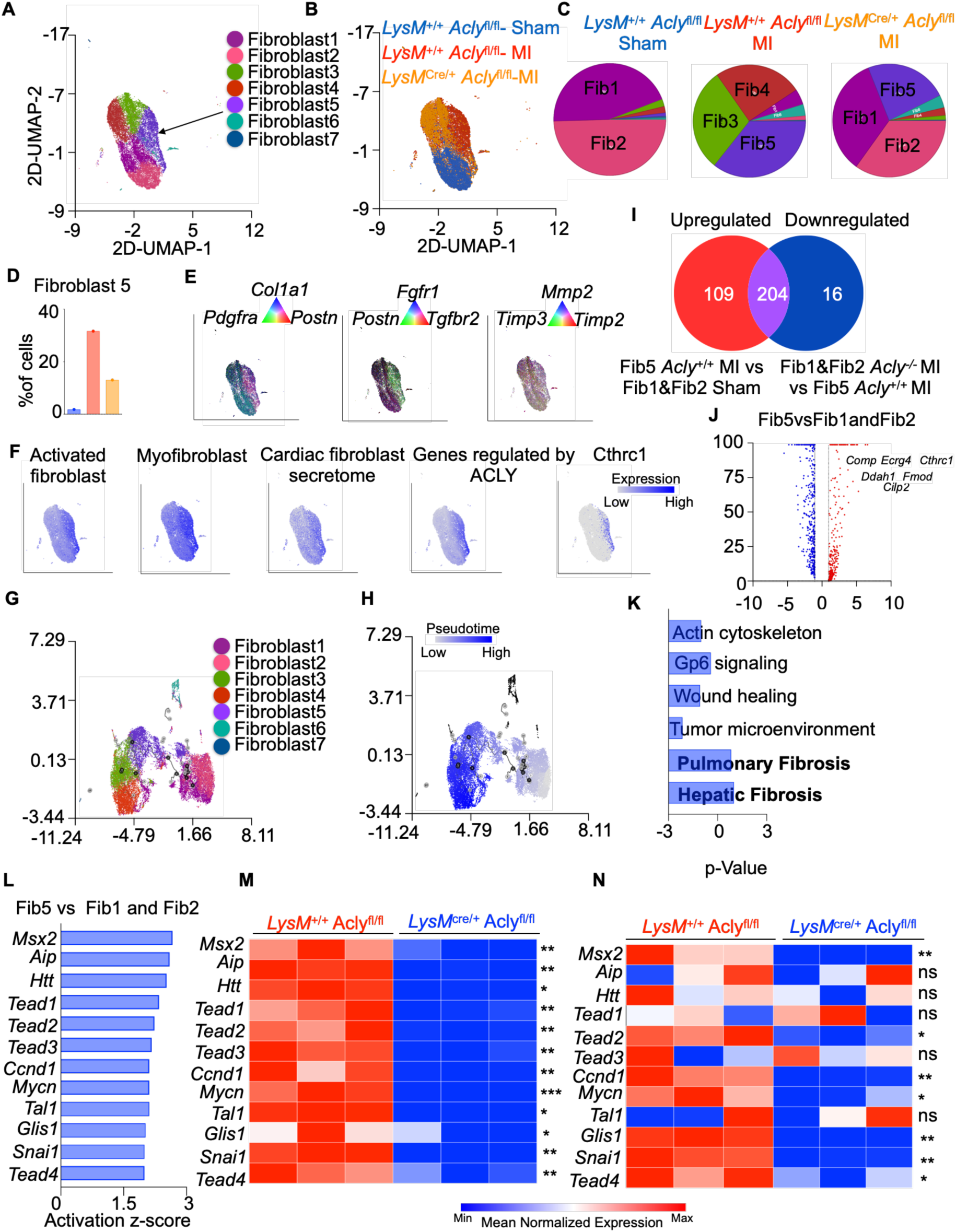
Macrophage ACLY promotes Fibroblast 5 expansion after MI. Cardiac fibroblasts were isolated from the heart of *LysM*^+/+^ *Acly*^fl/fl^ mice with sham and MI surgeries and *LysM*^Cre/+^ *Acly*^fl/fl^ mice with MI on day 28 after the surgery. Single cell RNA sequencing was performed on the sorted cells. n=4/ group. **A.** Uniform manifold approximation and projection (UMAP) plot depicts different fibroblast subtypes **B-C.** The UMAP plot (B) and pie charts (C) display the distribution of the fibroblast subsets. (**B**). **D.** The proportions of Fibroblast 5 are displayed. **E.** The UMAP plots and numeric triads show the expression of the genes expressed by activated fibroblasts. **F.** The expression of the gene sets reported to be expressed by activated fibroblast and myofibroblast, genes encoding cardiac fibroblast secretome, genes regulated by ACLY, and Cthrc1 is shown. **G-H.** Monocle3 was used to generate the Pseudotime trajectory plot of the fibroblast subsets. The trace from right to left reveals the transformation of resting fibroblasts to myofibroblasts. **I.** The Venn diagram reveals 204 common genes in 1. upregulated genes in Fibroblast 5 in *Acly*^+/+^ mice after MI compared to Fibroblasts 1&2 in mice with the sham surgery and 2. downregulated genes in Fibroblasts1&2 in *Acly^−/−^* mice with MI compared to Fibroblast 5 in *Acly*^+/+^ mice with MI. **J.** The volcano plot reveals the differentially expressed genes in Fibroblast 5 compared to Fibroblasts 1 & 2. **K.** A pathway analysis using the differentially expressed genes in J reveals the biological pathways enriched in Fibroblast 5. **L.** The bar graph lists the transcription factors regulating the differentially expressed genes in Fibroblast 5. **M-N.** The expression of the transcription factors in in cardiac fibroblasts of *LysM^+/+^ Acly*^fl/fl^ and *LysM^cre/+^ Acly*^fl/fl^ mice with MI (**M**) and C57BL/6 cardiac fibroblasts cultured in presence of *LysM^+/+^ Acly*^fl/fl^ and *LysM^cre/+^ Acly*^fl/fl^ BMDM (**N**). The qPCR data are expressed as mean ± SEM and pooled from 3–5 independent experiments. The Mann-Whitney U test was performed to test the statistical significance. \**p* < 0.05, \*\**p* < 0.01, \*\*\**p* < 0.001.

### ACLY^high^ cardiac macrophages are closer to Fibroblast 5-like cells in the heart of patients with MI

We next evaluated the presence of ACLY^high^ macrophages in human hearts and examined their distribution in relation to Fibroblast 5-like cells. To this end, we analyzed publicly available datasets of fibrotic human hearts from patients with MI^46^. As expected, single nuclear RNA seq demonstrated that the fibrotic zone was enriched for the myofibroblast genes compared to the control hearts (**Figure 7A**). Interestingly, the expression of Fibroblast 5 genes was elevated in the fibrotic zone (**Figure 7B**). Single nuclear ATAC seq revealed similar differences in summed peak intensities in these two patient groups (**Figures 7C-D**). To delineate relative distribution of cardiac macrophages expressing high or low levels of *ACLY* and cardiac fibroblasts expressing the Fibroblast 5 gene signature (**Table S4**) at high (Fib5^high^) or low (Fib5^low^) levels, we analyzed spatial transcriptomics data in the human heart. *ACLY*^high^ macrophages were closer to Fib5^high^ cells than *ACLY*^low^ macrophages (**Figures 7E&G**) whereas we did not find this spatial distribution pattern of these two subsets of cardiac macrophages in relation to myofibroblast location (**Figures 7F&H**). Additionally, the frequencies of Fib5^high^ cells (**Figure 7I**) but not myofibroblasts (**Figure 7J**) were higher near to *ACLY*^high^ cardiac macrophages compared to *ACLY*^low^ cardiac macrophages at nearer neighboring locations (≤12) from these macrophages. In summary, these results demonstrate the presence of a fibroblast population expressing the Fibroblast 5 gene signature and their proximity to ACLY^high^ macrophages in the hearts of patients with MI.

**Figure 7:**
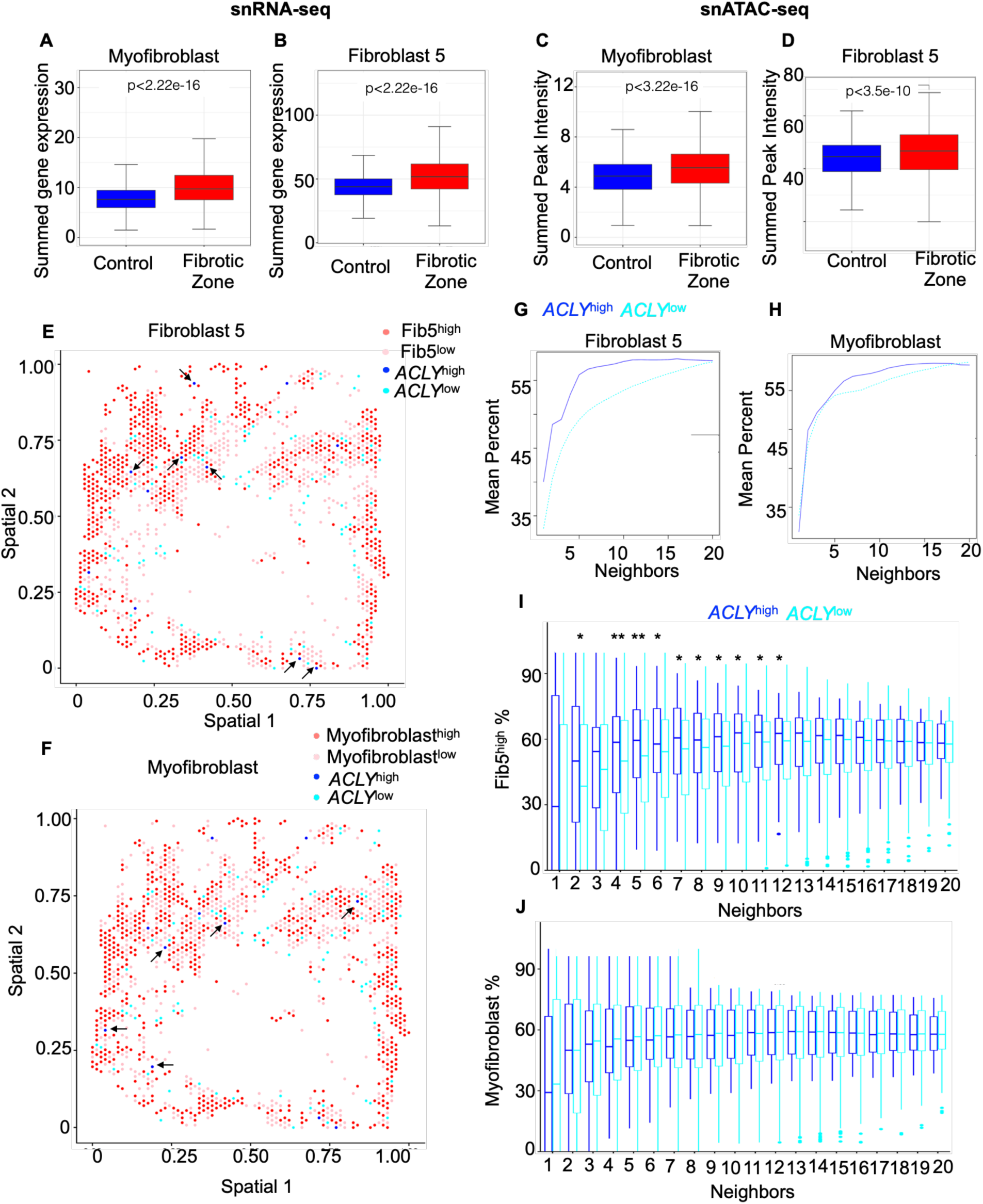
Fibroblast 5 expands in the heart in humans with MI. Publicly available single nuclear RNA and ATAC seq and spatial transcriptomics data in the heart of patients with MI were analyzed to detect cardiac macrophages expressing ACLY and myofibroblasts having the Fibroblast 5 gene signature. **A-D.** The box plots reveal summed expression (**A&B**, single nuclear RNA seq data) and peak intensity (**C&D**, single nuclear ATAC seq data) of the genes specific to myofibroblasts and Fibroblast 5. **E-F.** Spatial transcriptomics data analysis unveils the relative distribution of fibroblasts expressing high and low levels of the Fibroblast 5 genes (**E**) and fibroblasts expressing high and low levels of the myofibroblast genes (**F**) relative to macrophages with high and low levels of *ACLY* in the heart. **G-H**. The data displayed in E (**G**) and F (**H**) are summarized. **I-J** The box plots display the frequency of Fibroblast 5^high^ (**I**) and myofibroblasts (**J**) near *ACLY*^high^ or *ACLY*^low^ macrophages. The data are expressed as mean ± SEM. \**p* < 0.05, \*\**p* < 0.01.

## Discussion

Fatty acids can mediate inflammation in various ways-changing the lipid composition of the cell membrane and directly acting on cell surface or intracellular receptors^47^. Moreover, inflammatory macrophages synthesize fatty acids, which are used as precursors to produce inflammatory cytokines^9^. Impairment of fatty acid synthesis in the absence of FASN leads to aberrant composition of the cell membrane and defective trafficking of Rho GTPase required for inflammatory signaling^48^. Unexpectedly, FASN not only promotes the production of fatty acids but also cholesterol, which is required for TLR signal transduction^49^.

We show that cardiac macrophages synthesize *de novo* fatty acids after MI. There are concomitant increases in fatty acid synthesis enzymes in cardiac macrophages residing in the infarct. These enzymes are crucial in unleashing inflammation, precipitating cardiac fibrosis, and inciting cardiac remodeling. Nevertheless, our study does not ascertain whether newly made fatty acids magnify inflammation in macrophages.

The degree of cellular inflammation as measured by patients’ peripheral white blood cell counts has a positive correlation with in-hospital mortality following MI^50^, suggesting that systemic inflammation is crucial for MI prognosis. Consistently, in a recent clinical trial, IL-1β neutralization dramatically decreased major cardiovascular mortality and reinfarction in individuals following MI^51^. Previous research by us and others has demonstrated that elevated inflammation in atherosclerotic plaques drives reinfarction^12, 13^. These findings indicate that inflammation plays a major role in the outcome of MI. Yet, none of the current treatments for patients with MI are designed to suppress inflammation.

Bempedoic acid, an ACLY inhibitor, has been shown to help lower LDL cholesterol levels in patients in a recent study^52^. In patients with elevated lipid levels, *ACLY* genetic variations that inhibit gene activation reduce cardiovascular risk in addition to lowering LDL cholesterol levels^53^. Moreover, targeting macrophage *Acly* stabilizes atherosclerotic plaques^54^ with increased collagen deposition and fibrous cap thickness. A recent elegant study showed that ACLY modulates the myofibroblast gene program by acetylating histones^55^. Thus, ACLY inhibition is a potential therapeutic approach in cardiovascular disease. In another study, activation of FASN in stressed myocardium was found to be a compensatory reaction to shield cardiomyocytes from abnormal calcium flux^56^. Consistently, we observed a significant augmentation of FASN and ACLY post MI. Our preclinical study revealed the therapeutic benefit of targeting *Acly* in macrophages to ameliorate MI complications. Well-controlled clinical trials with bempedoic acid, an FDA-approved ACLY inhibitor, in MI patients are required to discern the therapeutic efficacy of this approach. Furthermore, in line with the reports on cardioprotective function of anti-inflammatory therapy, our study uncovered attenuated inflammation in mice with myeloid *Acly* deficiency. Congruently, ACLY inhibition thwarts the production of pro-inflammatory cytokines in LPS-stimulated macrophages^57^ and provokes the emergence of pro-reparative macrophages via Akt-mTORC1 signaling^58^.

As discussed above, ACLY cleaves citrate into acetyl-CoA, necessary for histone acetylation, which allows transcription factors to access chromatin structure. Histone acetylation supported by ACLY regulates the functions of a variety of cell types including adipocytes^22 59^, immortalized fibroblasts^22^, skeletal muscle^60^, and multiple cancer cell lines^61 62^. Moreover, the acetylation of the NF-κB promoter by ACLY increases the expression levels of proinflammatory genes in human macrophages^63^. We observed an increase in histone acetylation in pro-fibrotic genes like *Il33* ^27^ and *Nlrp3* ^64^ in macrophages in the presence of ACLY. Furthermore, we found that ACLY regulated the acetylation and expression of Krt17, which regulated the production of pro-fibrotic cytokines such as CCL17, CCL22, CCL24, CXCL2, and IL10, and induced fibrosis after MI. Because we were not able to retrieve sufficient cardiac macrophages to study the changes in their epigenetic marks, we elucidated the significance of ACLY in H3K27ac in BMDM. Although BMDM are primary macrophages, their functions are different from cardiac macrophages. New techniques should be developed to evaluate the epigenetic modifications in cardiac macrophages after MI.

After an ischemic injury, activated cardiac fibroblasts proliferate and repair the myocardium by producing extracellular matrix, which initiates fibrosis. Newly generated myofibroblasts are derived from resident fibroblasts in cardiac inflammation^65, 66, 67^. Mouse heart harbors several subsets of fibroblast populations^68^. However, after chronic angiotensin II infusion^69^ and MI^70^, specific subsets of fibroblasts emerge by selective proliferation and promote fibrosis. In congruence with the studies, our single cell RNA seq data in the chronic phase of remodeling post-MI showed expansion of novel cardiac fibroblast populations. We observed that Fibroblast 5 was enriched for genes promoting extracellular matrix synthesis, processing, and deposition. We also detected a fibroblast population expressing the Fibroblast 5 gene signature in the heart of MI patients and these fibroblasts but not myofibroblasts were located near *ACLY*^high^ cardiac macrophages, suggesting a crosstalk between this macrophage subset and Fibroblast 5-like cells. In conclusion, our study indicates that macrophage ACLY and FASN promotes cardiac remodeling after MI by expanding pathogenic cardiac fibroblasts.

## Materials and Methods

### Approval of the study

The University of Pittsburgh’s Institutional Animal Care and Use Committee (IACUC) provided its approval to conduct all animal experiments for this study. The University of Pittsburgh’s Institutional Review Board (IRB) approved the human studies performed in this project (STUDY19090164).

### Human studies

The left ventricles from patients with and without MI were obtained from National disease research interchange (NDRI). Heart tissues were embedded in OCT blocks and stored at −80° C. Tissue was sectioned using a cryostat and stained for immunofluorescence microscopy as discussed below. Additionally, we have analyzed publicly available data in patients with MI^46^.

### Experimental animals

All mice were bred on the C57BL/6 background. Deletion of *Acly* (The Jackson lab, #030772) and *Fasn* (Taconic Biosciences, #8034) in myeloid lineages were achieved by crossing these floxed transgenic mice with *Lysm*^Cre/Cre^ mice (The Jackson lab, #004781). After the breeding for three generations, mice with the genotypes *LysM*^+/+^ *Acly^fl^*^/fl^ or *LysM*^+/+^ *Fasn^fl^*^/fl^ were used as controls, and the mice with the genotypes *LysM^cre/+^ Acly*^fl/fl^ or *LysM^cre/+^ Fasn*^fl/fl^ had deficiency of the genes in myeloid cells. The experiments were performed in 10–12 week-old mice. The mice were housed in pathogen-free conditions and provided with food and water. Ambient temperature of 18 to 23° C, 40 to 60% humidity, and 12 hours light and dark cycles were maintained for housing these experimental mice. The following primers were used for genotyping: *Acly*^fl/fl^: Forward-CCCTCAGAAGGTCAGAGAACA and Reverse-CAGCAGGAGAGCTAGGACCA, *Fasn*^fl/fl^: Forward-GTTATCTACAGTTTCGACCCCACAG, Reverse-GCCAATGTTGTTGGTGAAATGACAAATGTCC, *LysM*^Cre^: Mutant-CCCAGAAATGCCAGATTACG, Common-CTTGGGCTGCCAGAATTTCTC, and Wild type-TTACAGTCGGCCAGGCTGAC.

### Coronary artery ligation

Coronary artery ligation was performed as described before ^16, 71^ In short, mice were injected with 0.05 mg/kg body weight of buprenorphine SR before the surgery. The mice were anesthetized with 2% isoflurane. Using toe pinch, the anesthetic depth was determined. After shaving the fur on the left thorax, the mice were intubated and kept on a heating pad at 37°C. The surgical area was cleaned with ethanol and betadine. Before the thoracotomy at the fourth left intercostal gap, the mice were placed on a ventilator. The pericardium was excised, the left anterior descending coronary artery was located, and it was ligated using a monofilament 8-0 nylon suture. A 5-0 suture was used to seal the thorax, skeletal muscle, and the skin. Daily wound monitoring was conducted to ensure timely recovery following surgery.

### D_2_O Treatment

For MALDI analysis, C57BL/6 mice having sham and MI surgeries were provided with deuterated water D_2_O (30% v/v) (Themo Fisher Scientific, 166301000) in drinking water for 28 days. As control, another group of mice with sham and MI surgeries received water (H20) for 28 days. The mice were euthanized, and organs were collected for MALDI analysis.

### DOTAP injection for gene silencing

Mice (22-25g) were injected with 33 µg of DOTAP (Encapsula NanoSciences, GEN-700) to deliver *Krt17* siRNA. At first, siRNA and DOTAP at the ratio of 1:7.5 were mixed, and the mixture volume was adjusted to 100 µl by adding RNAase and DNAase-free PBS. The mixture was incubated on ice for 15-20 mins and injected i.v in mice. The injections were performed immediately after MI twice a week for 28 days.

### IL-33 injection

MI was induced in *LysM*^+/+^ *Acly*^fl/fl^ and *LysM*^Cre/+^ *Acly*^fl/fl^ mice. About half of the *LysM*^Cre/+^ *Acly*^fl/fl^ mice with MI were injected with recombinant IL-33 (2 μg) (Biolegend, 580508) i.p every day for 28 days as described previously^72^. Rest of the mice were injected with the vehicle.

### Cardiac Magnetic resonance imaging (cMRI)

Cardiac function was evaluated with cardiac MRI ^73^. Briefly, mice on day 28 after MI were placed on a mouse stand in a supine position. A mixture of 2-3% isoflurane and oxygen was used to induce anesthesia. A constant temperature of 33 ± 2 °C was maintained to guarantee a sufficient and steady heart rate. Cine pictures of the short axis of the left ventricle were obtained using a 7 Tesla horizontal bore Pharmscan (Bruker) and a custom-built mouse cardiac coil in birdcage design (Rapid Biomedical). Based on ECG and respiratory signal, fast low angle shot sequences were obtained. We used the following parameters-2.7 ms for the echo time, 30 degrees flip angle, and 60 degrees for delayed enhancement imaging. The software Segment was employed for image analysis and quantification.

### Extracellular volume (ECV)

Myocardial fibrosis was assessed by measuring the extracellular volume (ECV) using gadolinium (Gd) contrast MRI. Subcutaneous injection of Gadobenate dimeglumine, (529 mg/ml), was administered via a sterile 24 Gauge subcutaneous catheter with the dosage of in 0.1 mmol Gd/kg bodyweight. Quantitative spin-lattice relaxation time (T_1_) was quantified by varied flip angle (VFA) method ^74, 75, 76^. Quantitative T1 mapping was acquired twice before and after administration of Gd contrast agent with the following parameters: Field of view (FOV) = 2.5 cm X 2.5 cm, slice thickness = 1 mm, in-plane resolution = 0.98, flip angles (FA) = 2, 19, 22, 28 degrees, echo time (TE) = 3.059 msec, repetition time (TR) = 5.653 msec, total scan time = 3 min 2 sec. Quantitative voxel-wise T1 maps were generated with custom-built MATLAB (The MathWorks Inc. Natick, MA). Hematocrit was measured from a droplet of blood from the facial vein. Quantitative voxel-wise ECV maps were generated with the established method ^77, 78, 79, 80^. Myocardial ECV = (1 − hematocrit) × (ΔR1_myocardium_/ΔR1_blood_), where R1 = 1/T1.

### Myofiber Architecture by Diffusion Tensor Imaging (DTI)

Myofiber architecture of the heart was evaluated with diffusion tractography derived from diffusion tensor imaging (DTI) using our established method ^81, 82^

#### Sample Preparation

After the mouse was euthanized, the blood in its heart was removed by retrogradely perfusing it with PBS through the abdominal aorta for two minutes. Following that, the tissue was fixed by retrogradely perfusing it with 4% paraformaldehyde (PFA). The fixed mouse heart was taken out of the formalin for the MRI and gently patted dry with paper towels to eliminate any extra water from the heart’s surface. The hearts were then transferred to 1.5 cc tube containing Fomblin-Y perfluoropolyether vacuum oil (Millipore Sigma, MW = 1800), to remove the susceptibility artifact of the tissue-to-air interface.

#### DTI Acquisition and Myofiber Tractography

Using a DtiStandard SpinEcho procedure, the diffusion pictures were obtained on a Bruker BioSpin MRI GmbH 7-Tesla scanner. TE=16.945 msec, and TR=900 msec. The diffusion time was 8 msec. The diffusion encoding duration was 4 msec. Thirty diffusion sampling directions were obtained using a DTI diffusion method. The b-value was 1044.64 s/mm2. The in-plane resolution was 0.15625 mm. The slice thickness was 0.127907 mm. The diffusion data were reconstructed using generalized q-sampling imaging ^83^ provided by the DSI studio (https://dsi-studio.labsolver.org/) with a diffusion sampling length ratio of 1.25, angular threshold 60°, the step size 0.021 mm, the anisotropy threshold 0.0263, and total 1,000,000 tracts calculated. The restricted diffusion was quantified using our restricted diffusion imaging protocol ^82^.

### Echocardiography

Echocardiographic analysis was performed on conscious mice after sham or MI surgery using the Vevo 3100 ultrasound machine with the MX400 linear transducer capable of 40 MHz (Visualsonics). The rodent is anesthetized using 3% isoflurane mixed with 1 L/min of 100% oxygen in an induction chamber. Once anesthetized, the rodent is then transferred to a warming table and positioned for imaging using 1.5% isoflurane. The heart rate was maintained between 400-500 bpm during ultrasound imaging. Both B-Mode and M-Mode imaging was performed. The parasternal long axis (PS-LAX) view was obtained, and measurements were taken by area method using the area of the left ventricle (LV) in diastole and systole. The parasternal short axis (PS-SAX) was imaged and acquired at mid papillary muscle. All imaging was performed by a blinded sonographer. The data were analyzed on B mode PS-LAX and M mode PS-SAX. The LV was traced throughout the complete cardiac cycle. In B mode, PS-LAX full diastole was located starting at the R wave. Two points were drawn at the aortic roots, one point at the apex and other two points on both sides of the LV. The next appropriate full systole frame was navigated with a combination of eyes and the physiotrace (ECG) and was traced like previously. The smallest trace was highlighted in green, and the largest trace (diastole) was highlighted in red. The ejection fraction (EF), stroke volume (SV), cardiac output (CO), and diastolic (dsVol), and systolic volume (sVol) were obtained by the method of LV tracing as described. Fractional shortening (FS) and Left ventricle mass (LV-mass) was obtained by M mode PS-SAX data analysis by tracing the LV during systolic and diastolic phases.

### Organ harvesting and flow cytometry

Following MI, peripheral blood was collected from mice using the retroorbital bleeding technique in a tube with a 50 mM EDTA solution. (Invitrogen, AM9260G). A commercially available RBC lysis buffer (BioLegend, 420301) was used to lyse the RBCs in the blood. Next, the left ventricle was perfused by administering PBS to remove blood from organs. The infarct, border, and remote areas of the heart were excised, minced, and processed into single-cell suspension in a mixture of 450 U/mL collagenase I (Gibco, 17100017), 125 U/mL collagenase XI (Sigma-Aldrich, C7657), 60 U/mL DNase I (Qiagen, 79254), and 60 U/mL hyaluronidase (Sigma-Aldrich, H35006) at 37°C for 1 hour. Following digestion, the tissue was passed through a 70 μm cell strainer, and the cells were resuspended in FACS buffer (0.5% BSA in PBS). After removing the spleen and mincing it in FACS buffer, the cell solution was passed through a 70 μm cell strainer. The femur, tibia, and fibula were flushed with FACS buffer to prepare single-cell suspensions. The cell suspensions from the heart, blood, bone marrow and spleen were counted using a hemocytometer and stained to enumerate Leukocytes (CD45^+^) myeloid cells (CD45^+^ CD11b^+^), monocytes (CD45^+^ CD11b^+^ CD115^+^ Ly-6G^−^), macrophages (CD45^+^ CD11b^+^ CD115^−^ Ly-6G^−^ F4/80^+^), neutrophils (CD45^+^ CD11b^+^ CD115^−^ Ly-6G^+^), T cells (CD45^+^ CD11b^−^ CD3^+^), B cells (CD45^+^ CD11b^−^ CD19^+^), and dendritic cells (CD45^+^ CD3^−^ CD11c^+^) using the following antibody F4/80 PE-Cy7 (Biolegend,123114), CD11b APC-Cy7 (BD Biosciences, 557657), CD45.2 Alexa fluor 700 (BioLegend, 109822), CD3 BV 421 (BD, 564008), CD19 BV 605 (BD, 563148), Ly6G APC (Biolegend, 127614), CD115 PE (Biolegend, 135506) and CD11c BV 510 (BioLegend, 117338) by Cytek Aurora flow cytometer (Cytek Biosciences). Fluorochrome compensation was performed using beads (Invitrogen, 01-3333-42). Data were analyzed using FlowJo X 10.4.1 (Tree Star Inc.). Viability dyes were used to discriminate between living and dead cells.

### Isolation, culture, and treatment of murine bone marrow derived macrophage (BMDM)

The leg bones, comprising the tibias, fibulas, and femurs, were removed following euthanasia. The bones were flushed with DMEM supplemented with 4.5 g/L glucose (Gibco, 11965118), 20% L-929 conditioned medium, 10% FBS (VWR, 10803-034) and 1% penicillin/streptomycin (Gibco, 15140122). Following four days of culturing in 100 mm dishes, the old cell culture media were replaced with media. Cells were allowed to differentiate for 7 to 10 days with changing of the media every 2 to 3 days. ^84^. The cells were plated at a confluency of 1× 10^6^ cells/well or 0.5× 10^6^ cells/well, respectively, in 6-well or 12-well plates prior to various treatments.

### siRNA transfection *in vitro*

Small interfering RNAs (siRNAs) targeting *Ikbkb*, *Nlrp3*, *Egr1*, *Krt17*, *Ikbkg*, and *Chuk* were transfected into BMDMs at a concentration of 25 nM using Lipofectamine 3000 Transfection Reagent (Invitrogen, L3000008) in accordance with the manufacturer’s instructions for gene-silencing studies. With RT-qPCR, the knockdown efficiency was assessed.

### Cardiac fibroblasts isolation and culture

Adult mouse fibroblasts were separated by removing the hearts from euthanized mice. Hearts were minced in 10 ml of digestion buffer containing 100U/ml of collagenase II (Gibco, 17101015) and 0.1% Trypsin (Gibco, 15090046) in HBSS (Gibco, 14175095). The minced heart tissue was transferred into sterile falcon tubes and digested in a cell culture incubator for 15-20 minutes. The samples were passed through cell 70 μm cell strainers. The cell suspensions were plated in 60 mm Petri dishes and incubated in cell culture incubator for 4 hours. The supernatants were then discarded, and adhered fibroblasts were rinsed with warm sterile 1X PBS. The adhered cells were then cultured in a humidified cell culture incubator at 37°C and 5% CO_2_ in fibroblast medium containing DMEM/F12 (Gibco, 11320033), 10% FBS, 100 U/ml PenStrep, 1x L-glutamine (Gibco, 25030081), and 100 µM ascorbic acid (Thermoscientific, A15613.22) acid for a week. Cardiac fibroblasts were then used for different experiments. Cardiac fibroblasts were grown for 3 to 5 passages before using them for experiments. The purity of cultured cardiac fibroblasts was verified using the fibroblasts markers MEFSK4-APC (Milteny Biotec, 130-120-802) and Vimentin (Abcam, ab92547)^85^.

### Coculture of BMDM and cardiac fibroblasts

BMDM isolated from WT or *Acly* KO mice and cardiac fibroblasts isolated from C57BL/6 mice were cultured according to the protocol described above. These cells were detached using 0.25% Trypsin (Thermoscientific, 25200056) and washed with 1×PBS. These cells were then combined in 1:1 ratio in 10% FBS-supplemented DMEM at 37°C in a humidified environment containing 5% CO_2_. Cocultured cells were incubated at the above-mentioned conditions for 48 hours, and cardiac fibroblasts were sorted using FACs Aria.

### Proliferation assay

Cardiac fibroblasts cocultured with WT and *Acly* KO BMDM were assessed for proliferation. Briefly, after cell surface staining using the following antibodies CD45 AF 700, CD11b PECy7, and MEFSK4 APC, cells were fixed and permeabilized using the fixation (BD biosciences, 554655) and permeabilization (BD biosciences, 558050) buffers, respectively. The cells were then incubated with Ki67 BV605 antibody (BD biosciences, 567122) for intracellular antigen detection and incubated for 30 mins at RT protected from light. The cells were then washed using FACS buffer and analyzed with a flow cytometer (BD Fortessa). Cardiac fibroblasts in culture with BMDM were identified as CD45^−^ CD11b^−^ MEFSK4^+^, and their cell cycle status was assessed as G0-Ki-67^−^DAPI^−^, G1-Ki-67^+^ DAPI^−^, and S-G2M-Ki-67^+^DAPI^+^.

### Fluorescence-activated cell sorting (FACS)

To sort cardiac macrophages, single cell suspensions of cardiac tissue were used as described above in the flow cytometry method. The cells were stained for macrophage markers (CD45^+^ CD11b^+^ CD115^−^ Ly-6G^−^ F4/80^+^). To sort cardiac fibroblasts, mouse hearts particularly infarct and the border zone were digested using the digestion buffer mentioned above. Cardiac fibroblasts cocultured with BMDM were treated with trypsin and washed. The cell suspensions of either digested hearts or cocultured cells were stained with the following antibody cocktail CD31 PE-Cy7 (Biolegend, 102418), CD45.2 Alexa Flour 700 (BioLegend, 109822), Ter119 PE (Biolegend,116208), and MEFSK4 APC (Milteny Biotec, 130-120-802). Calcein (Invitrogen, C1430) and DAPI (Milteny Biotec, 130-111-570) were used to identify live and dead cells, respectively. DAPI^−^ Calcein^+^ CD31^−^ Ter119^−^ CD45^−^ MEFSK4^+^ cells were deemed as cardiac fibroblasts^30^. The sorted cells were collected either in RNA extraction buffer for transcriptomics and gene expression analyses or in FACS buffer for immunofluorescence.

### Immunomagnetic cell separation

Cardiac fibroblasts from WT and KO mice having sham and MI surgeries were sorted by negative and positive selection using EasySep™ Biotin Positive Selection Kit II (Stem cell technologies, 17683). For negative selection, biotinylated CD45.2 (Biolegend, 109803), CD31(Biolegend,102503), and Ter119 (Biolegend, 116204) were used to label the single cell suspensions of cardiac tissues. The mixture of the biotinylated antibodies and cell suspensions were incubated on ice for 30 minutes. The cell suspensions were then washed and incubated with 100 µl of biotin selection cocktail for 15 minutes at room temperature. To this mixture, 50 µl of magnetic nanoparticles were added and for 10 minutes in room temperature. The total volume of the mixture was adjusted to 2.5 ml in a FACS tube and placed in a EasySep Magnet (Stem cell technologies,18000). The beads were discarded, and the unbound cells were incubated with biotinylated MEFSK4 antibody (Milteny Biotec, 130-102-015) for positive selection. After washing and incubating with magnetic nanoparticles, the cells were placed in a EasySep Magnet. The cells bound to the magnetic beads were collected, washed, and assessed for viability.

### Bulk RNA seq analysis

FACS Aria was employed to sort cardiac macrophages into RNAse-free tubes containing 100 µl of RNA extraction buffer. A commercially available PicoPure RNA isolation kit (Applied Biosystems, #12204-01) was used to extract mRNA from the cells. The Health Science Sequencing Core at UPMC Children’s Hospital of Pittsburgh carried out the library preparation and mRNA sequencing, and FASTQ files were submitted to GSE236572. RNA sequencing data analysis was carried out using Qiagen CLC Genomics Workbench 21, and a quality control (QC) report was produced to verify the quality of the data. From the 3’ end of the reads, the adaptor sequences were removed. Following trimming, the reads were mapped against the version of the reference genome of *Mus musculus* (mm10) in CLC genomics workbench. Reads Per kilobase per Million mapped reads (RPKM) was used to normalize gene expression, and the genes that met the cut off of FDR P<0.05 and >2 log2 fold change in Acly KO compared to WT cardiac macrophages were identified. To show differential gene expression (DEG) in KO vs WT, a volcano plot was created. A heatmap was generated to show the genes that met the above mentioned cut off. Next, we employed Ingenuity Pathway Analysis (IPA) to detect disease and classical pathways and upstream regulators of the genes that were expressed differentially in the KO vs WT cardiac macrophages.

#### scRNA-seq

After assessing cell viability of sorted cardiac fibroblasts from WT and *Acly* KO mice, about 7000 cells per sample were used for 10x Genomics library preparation for scRNA sequencing. The Partek Flow software (version 10.0.22.1204) was used to analyze the count matrix generated from the sequencing data. During analysis, cells were filtered applying these standards: read counts-15000, detected features-2500, mitochondrial reads-5%, and ribosomal counts-50%. Using counts per million and Log2 transformation, the overall UMI (Unique Molecular Identifier) counts were normalized according to the recommendations of Partek Flow. Following normalization, we further filtered the data by eliminating features for which at least 99 percent of the cells had gene expression values of ≤1. With the Louvain algorithm, the graph-based clustering analysis was carried out. To visualize the single-cell data, we performed a Uniform Manifold Approximation and Projection (UMAP) dimensional reduction and generated the plots. Cell based clustering was performed by identifying the cell types in PangolaoDB database using the top five markers of each cluster. Using Partek flow software, the distribution of transcriptomically different fibroblasts was observed in Sham, MI and AclyKO+MI group and represented as Pie-chart and bar graphs. The differentially expressed genes (DEGs) in pathogenic (Fib5) vs Sham (Fib1 and Fib2) fibroblasts were obtained using gene specific analysis (GSA) algorithm built in Partek flow software. The DEGs were filtered based on FDR P<0.05 and 2 log2 fold change. Heatmaps and volcano plots were constructed using this information. IPA analyses were performed using DEGs in pathogenic vs. sham fibroblasts to ascertain upstream regulators and canonical pathways. Monocle3 in Partekflow software was employed to perform pseudo time trajectory analysis. The single-cell RNA-seq data were deposited to GEO (GSE237583).

#### Spatial scRNA-seq and scATAC-seq analysis

Visium spatial scRNA-seq and scATAC-seq datasets of fibrotic and control (normal) human hearts were downloaded from^46^ (https://cellxgene.cziscience.com/collections/8191c283-0816-424b-9b61-c3e1d6258a77). Wu used two different gene sets to compare fibroblast populations in human hearts: “Fibroblast 5” (obtained from our analysis of mouse hearts) and “Myofibroblasts” ^86^. The summed gene expression (or peak intensity for scATAC-seq) of each of these four gene sets was calculated for every fibroblast cell, and the cells with summed expression higher than the 50 percentile were considered “marker high” fibroblast cells (e.g. Fib5^high^) whereas those with summed expression less than the 50 percentile were considered “marker low” fibroblast cells (e.g. Fib5^low^). Among myeloid cells, we defined macrophages as those expressing at least one of the macrophage markers (**Table S6**). These macrophages were further classified into *ACLY*^high^ macrophages if Acly was expressed, otherwise *ACLY*^low^ macrophages. For each macrophage subset, the ratio of marker high fibroblast cells to total fibroblast cells was computed within each fixed neighbour step and averaged across *ACLY*^high^ or *ACLY*^low^ macrophage cells.

### MALDI

MALDI was performed to detect fatty acids, such as oleate, stearate, linoleate palmitate, and arachidonate, in mouse hearts at the Mass Spectrometry Research Center at Vanderbilt University. Immediately after sham or MI surgery, mice were provided with D_2_O in drinking water. The mice were euthanized 28 days after MI and their hearts were fresh frozen. Each heart was sectioned in12 µm sections. MALDI matrix (9-aminoacridine, 9AA, (Sigma-Aldrich) was spray-coated onto the target slides containing cardiac section in an automated fashion using a TM Sprayer (HTX Imaging). 9-AA was made up as a 5 mg/ml solution in 90% methanol. Four passes were used with a nozzle temperature of 85 °C, a flow rate of 0.15 ml/min, 2-mm track spacing, and a stage velocity of 700 mm/min. Nitrogen was used as the nebulization gas and was set to 10 psi. Images were acquired on a 15T Fourier transform ion cyclotron resonance mass spectrometer (FT-ICR MS, Solarix, Bruker Daltonics) equipped with an Apollo II dual ion source and Smartbeam II 2kHz Nd:YAG laser that was frequency tripled to 355 nm. Data were collected in the negative ion mode with the laser operating at 2 kHz at 35 μm resolution. Tentative metabolite identifications were made by accurate mass, typically better than 1 ppm. The sections were then stained with macrophage marker CD68 antibody (eBioscience, 14-0681-82), and whole sections were imaged using a confocal microscope. MALDI images were superimposed with IF images. Macrophage rich areas (50 μm) were selected to measure fatty acid levels. MALDI images were analyzed with flexImaging software (Bruker), while average spectra in macrophage rich regions were exported to mMass for visualization of differences.

### Immunofluorescence

Paraformaldehyde-fixed and OCT-embedded heart sections were cryosectioned into 12 µm sections. Triton X-100 (0.1%) and SDS (4%) were used for antigen retrieval and permeabilization by incubating cardiac tissue sections for 4 mins at room temperature. The sections and cells were blocked with 5% BSA/PBS for 60 min and stained overnight at 4 °C with primary antibodies against ACLY (Invitrogen, PA5-29495), ACC (Thermofisher, MA5-15025), FASN (Abcam, ab22759), CD68 (eBioscience, 14-0681-82), Vimentin (Abcam, ab92547), *⍺*SMA (Thermofisher, PA5-18292), F4/80 (Thermofisher, MA1-91124), and Ki-67 (Thermofisher, 9129S). Next, secondary antibodies conjugated with Alexa Fluor 488 (Invitrogen, A-11001 A-11008), Alexa Fluor 594 (Invitrogen, A-11007 A-21205) or Alexa Fluor 647 (Invitrogen, A-31634 A-21447 A21247) was added in the primary antibody-stained sections and cells. The sections were stained and fixed with Vectashield mounting medium with DAPI (Vector Laboratories, H-1200). Confocal microscopy was conducted to capture images using a Nikon A1 Spectral Confocal. The ImageJ software was used for image analysis.

### Masson’s Trichrome Staining

Hearts were collected after perfusing with PBS, fixed using 4 % paraformaldehyde, and embedded in OCT (optimum cutting temperature) compound (Fisher Scientific, 23-730-571) Sections of 6 µm thickness were made and stained using Masson’s trichrome staining reagents (Fisher Scientific, 22110648) to access fibrous area. VS200 research slide scanner was used to capture images. Percentage of fibrous area was calculated by dividing the fibrous area with total area of the heart.

### Chromatin Immunoprecipitation (ChIP)

ChIP was performed in WT and *Acly* KO BMDM as previously described^87, 88^. Briefly, BMDM from these mice were fixed with 1% PFA (ThermoFisher Scientific, 28908) for 10 min at room temperature. The chromatin fragments of 200-500 base pairs were obtained by sonicating cells with a Bioruptor Pico (Diagenode, B01060010). The fragmented chromatin was incubated with Protein G Dynabeads (Invitrogen, 10004D) and one of the following antibodies: H3K27Ac (Invitrogen, MA5-23516) and rabbit IgG (Abcam, ab171870). Phenol and chloroform were used to extract genomic DNA from immunoprecipitated (IP) and non-immunoprecipitated (INPUT) samples. The UCSC Genome Browser was used to identify H3K27Ac binding motifs on *Ikbkb*, *Nlrp3*, *Egr1*, *Krt17*, *Ikbkg*, and *Chuk* gene promoters. NCBI Primer BLAST was used to design the sequences of the primers flanking the binding motifs. qPCR was performed to quantify precipitated binding motifs. The results were expressed as IP/INPUT.

### CUT&RUN Assay

BMDM was treated with 0.5 mg/mL oxLDL (ThermoFisher Scientific, L34357) for 24 hours at 37°C. ^89^ The chromatin profiling was performed using the CUT&RUN assay kit (Cell signaling technology, 86652) as per the manufacturer’s protocol. Briefly, 100,000 BMDM plated in 12-well plates were collected in Eppendorf tubes, washed, and incubated with activated Concanavalin A beads for five minutes under rotating conditions. The supernatant was removed using a magnetic stand, and the cells were then resuspended in antibody binding buffer containing 1X Spermidine, 1X PIC, and 0.01% digitonin. To this, 2:100 µl of anti-H3K27Ac antibody (Invitrogen, MA5-23516) was added and rotated at 4° C overnight. Next day, the cells were collected and treated with 1.5 µl of pA/G-MNase enzyme for 1 hour under rotating conditions at 4° C to facilitate enzyme binding. Next, the cells were treated with 3 µl of cold CaCl_2_ and incubated at 4° C for an hour to initiate enzyme activation for digestion and release of targeted chromatin. The genomic DNA was purified using DNA Purification Buffers and Spin Columns (Cell signaling technology, 14209) as per the manufacturer’s instructions. Sequencing libraries were prepared using CUT&RUN Library Prep Kit (EpiCypher, SKU: 14-1001). We performed the sequencing of extracted genomic DNA with the help of Health Science Sequencing Core, UPMC Children’s Hospital of Pittsburgh.

### CUT&RUN data-processing and analysis

The analysis of CUT&RUN DNA sequencing was performed by using the Partek Flow (version 10.0.22.1204) software. In brief, pre-alignment QA/QC was first performed and then adaptors were trimmed before BWA alignment to the Mus musculus GRCm39 (mm39) genome. Aligned reads were then checked for post-alignment QA/QC and filtered. Peak calling was then performed using Model-based Analysis of ChIP-Seq (MACS). Peak annotation, integration with Bulk RNA seq data, and visualization of the data were carried out using the inbuilt algorithm of the Partek Flow software.

### RT–qPCR

Total RNA was extracted from sorted or cultured cells using Qiagen RNA isolation kit (Qiagen, 74104) and Picopure RNA solation kit (ThermoFisher Scientific, KIT0204). cDNA was synthesized from 1 μg of RNA using a cDNA synthesis kit (Applied Biosystems, 4387406). RT– qPCR was performed with an Applied Biosystem machine using SYBR green (Applied Biosystems, A25742), and the results were represented as Ct values normalized to the housekeeping gene β actin.

### Statistical analysis

Data are represented as mean±SEM. Statistical significance between groups was assessed, and graphs were created using Prism. The Mann–Whitney U test was used to compare differences between two group, and the one-way ANOVA was used for data sets containing more than two groups. Results were considered as statistically significant when *P*<0.05.

## Data availability

The article and its online supplementary files contain all of the information.

## Acknowledgments

This research was funded by the National Institutes of Health (NIH) grants R01HL142629, R01HL142629-04W1, R01HL143967, R01AG069399 and R01DK129339, the American Heart Association (AHA) Transformational Project Award (19TPA34910142), the AHA Innovative Project Award (19IPLOI34760566), the ALA Innovation Project Award (IA-629694), and the AHA Innovative Project Award (23IPA1053549) to P. Dutta. S. Sadaf received funds from the AHA Postdoctoral Fellowship Award (23POST1029135). The Center for Biologic Imaging (CBI), University of Pittsburgh, used NIH-supported grants to perform confocal and intravital microscopy. Specifically, the NIH grant 1S10OD019973-01 was used to fund the confocal microscope. The bulk RNA seq data analysis was carried out using CLC Genomics Workbench software, which is licensed through the Molecular Biology Information Service of the Health Sciences Library System (HSLS), University of Pittsburgh. Partek Flow software, version 10.0.22.1204, licensed by the HSLS University of Pittsburgh, was used to analyze scRNA and CUT&RUN sequencing data. The University of Pittsburgh Center for Research Computing, RRID:SCR_022735 supplied the resources necessary to support this research, in part. Specifically, this work used the HTC cluster, which is supported by NIH award number S10OD028483. The BioRender software was used to create the visual abstract. We thank Life Science Editors for editing services.

## Author Contributions

SS were involved in designing and executing the experiments, data analysis and manuscript writing. SV was involved in *LysM^Cre/+^Acly^fl/fl^* mouse generation and helping with organ harvest. IJ was involved in analyzing the qPCR data and tissue sectioning. EJ helped with IV injections and Masson’s trichrome staining KK and SH assisted with ELISA and Masson’s Trichrome staining. AM helped in genotyping of mice. SC helped in teaching to run the software like Partek flow and CLC genomics to SS to analyze ScRNA seq and Cut and Run sequencing data. SO helped in spatial transcriptomics data analysis. PD was involved in designing experiments, analyzing date, writing the manuscript, and providing the fund to complete the study.

## Declaration of interests

The authors declare no competing interests.

---

**Supplemental Figure 1:**
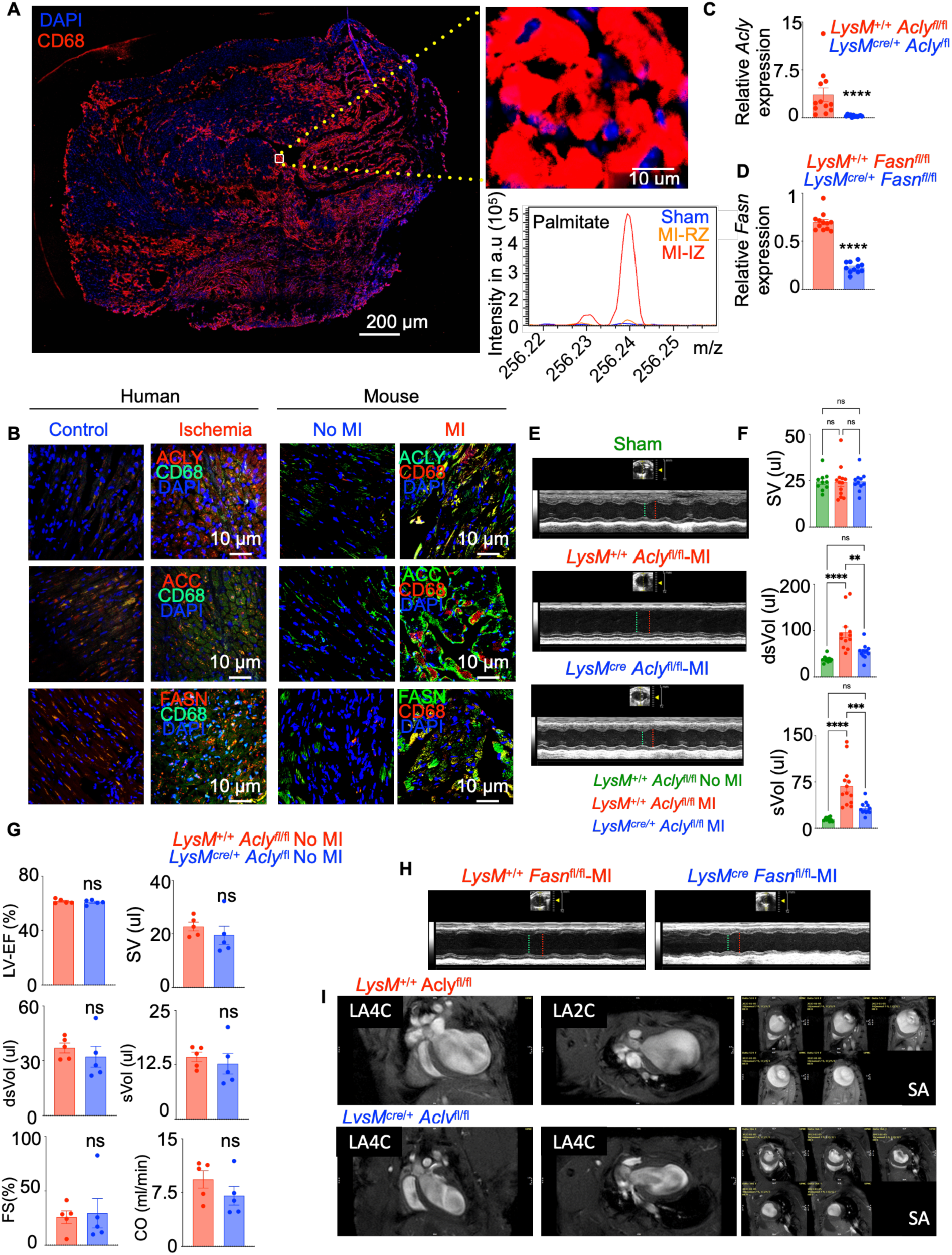
Myeloid *Acly* and *Fasn* deficiencies diminish cardiac remodeling after MI. **A.** MALDI and confocal microscopy were performed on the same cardiac sections in C57BL/6 mice provided with D_2_O in water 28 days after MI. Single spectra of palmitic acid in macrophage clusters were analyzed by mMass (n=3/group). **B.** ACLY, ACC, and FASN staining in macrophages (CD68+) is quantified by confocal microscopy in the heart of patients and mice with MI (n=3-5/group). **C-D.** qPCR quantification of *Acly* in *LyzM^+/+^ Acly^fl/fl^* and *LyzM^cre/+^ Acly^fl/fl^*(n=10-12/group) (**C**) and *Fasn* in *LyzM^+/+^ Fasn^fl/fl^* and *LyzM^cre/+^ Acly^fl/fl^* (**D**) cardiac macrophages (n=10-12/group). **E-F**. Representative echocardiogram M-mode images. The green and red lines indicate LVID;s = left ventricular internal diameter in systole and LVID;d = left ventricular internal diameter in diastole respectively (**E**) and quantification of heart function (**F**) in *LysM*^+/+^ *Acly*^fl/fl^ and *LysM^cre/+^ Acly*^fl/fl^ mice on 28 days after MI are provided (n=5-13/group). **G**. Echocardiography quantification in *LysM^+/+^ Acly^fl/fl^* and *LysM^cre/+^ Acly^fl/fl^* mice in the steady state-(without an MI) (n=5/group). **H**. The images depict representative echocardiograms of the LV in M-mode in SAX view in *LyzM^+/+^ Fasn^fl/fl^* and *LyzM^cre/+^ Fasn^fl/fl^* mice on day 28 after MI (n=6-10/group). The green and red lines indicate LVID;s = left ventricular internal diameter in systole and LVID;d = left ventricular internal diameter in diastole respectively **I.** Representative long axis four-chambered (LA-4C), long axis two-chambered (LA-2C), and short axis (SA) views of cine magnetic resonance imaging of *LysM*^+/+^ *Acly*^fl/fl^ and *LysM^cre/+^ Acly*^fl/fl^ mice on day 28 after MI (n=5/group) are shown. The data are expressed as mean ± SEM and pooled from 3–5 independent experiments. The Mann-Whitney U test was used for S1C-D and One way ANOVA was performed for Fig S1F \*\**p* < 0.01, \*\*\**p* < 0.001, \*\*\*\**p* < 0.0001.

**Supplemental Figure 2:**
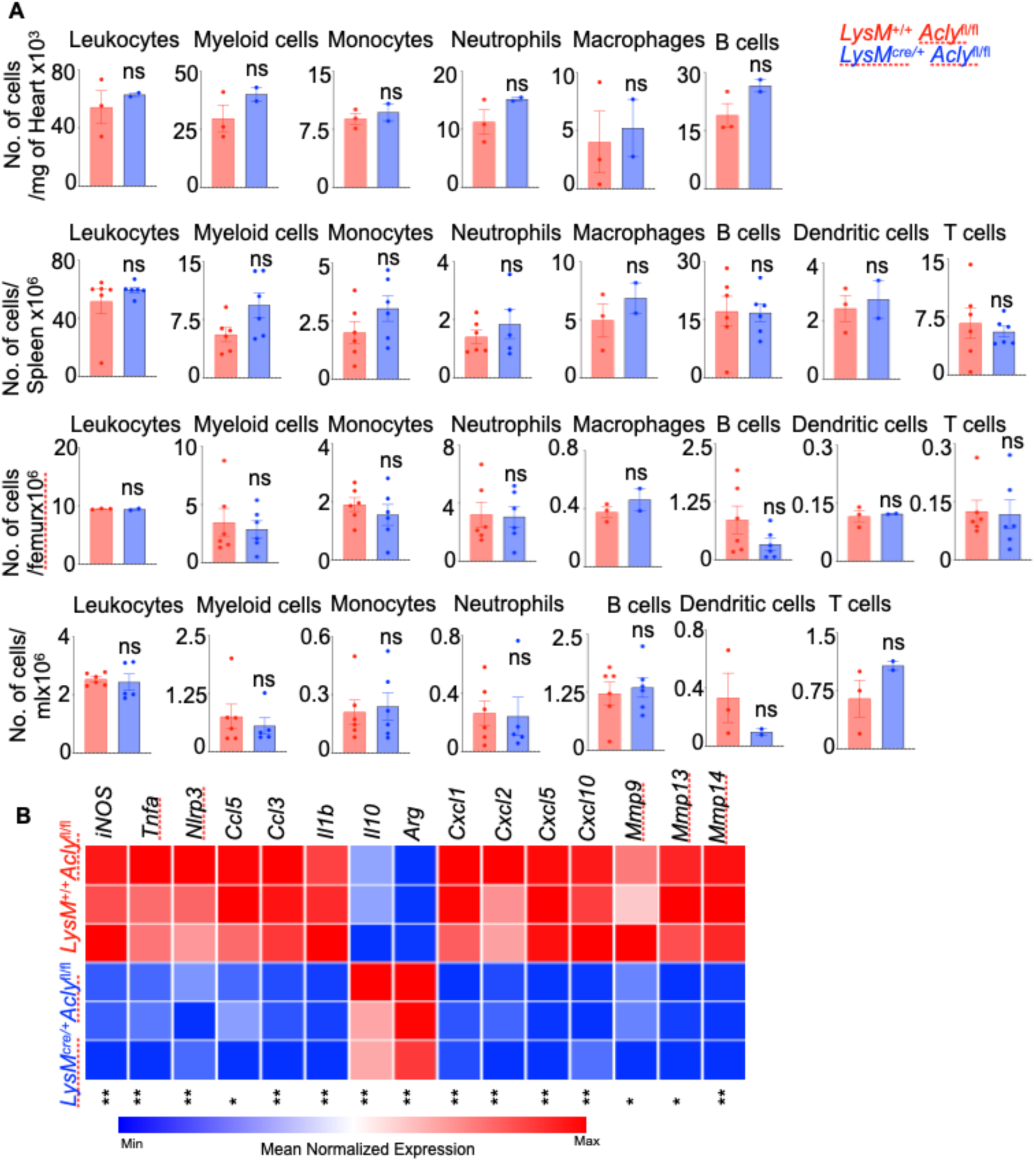
Myeloid *Acly* depletion suppresses inflammatory genes in the infarct after MI. **A.** Flow cytometry was employed to enumerate different leukocyte subsets in the heart, spleen, bone marrow, and blood in *LysM*^+/+^ *Acly*^fl/fl^ and *LysM^cre/+^ Acly*^fl/fl^ 28 days after MI (n=5-6/group). **B.** The heatmap displays inflammatory genes expression in cardiac macrophages sorted from the infarct of *LysM*^+/+^ *Acly*^fl/fl^ and *LysM^cre/+^ Acly*^fl/f^ mice on day 28 after MI (n=10-12/group). The data are expressed as mean ± SEM and pooled from 3–5 independent experiments. The Mann-Whitney U test was performed \**p* < 0.05, \*\**p* < 0.01.

**Supplemental Figure 3:**
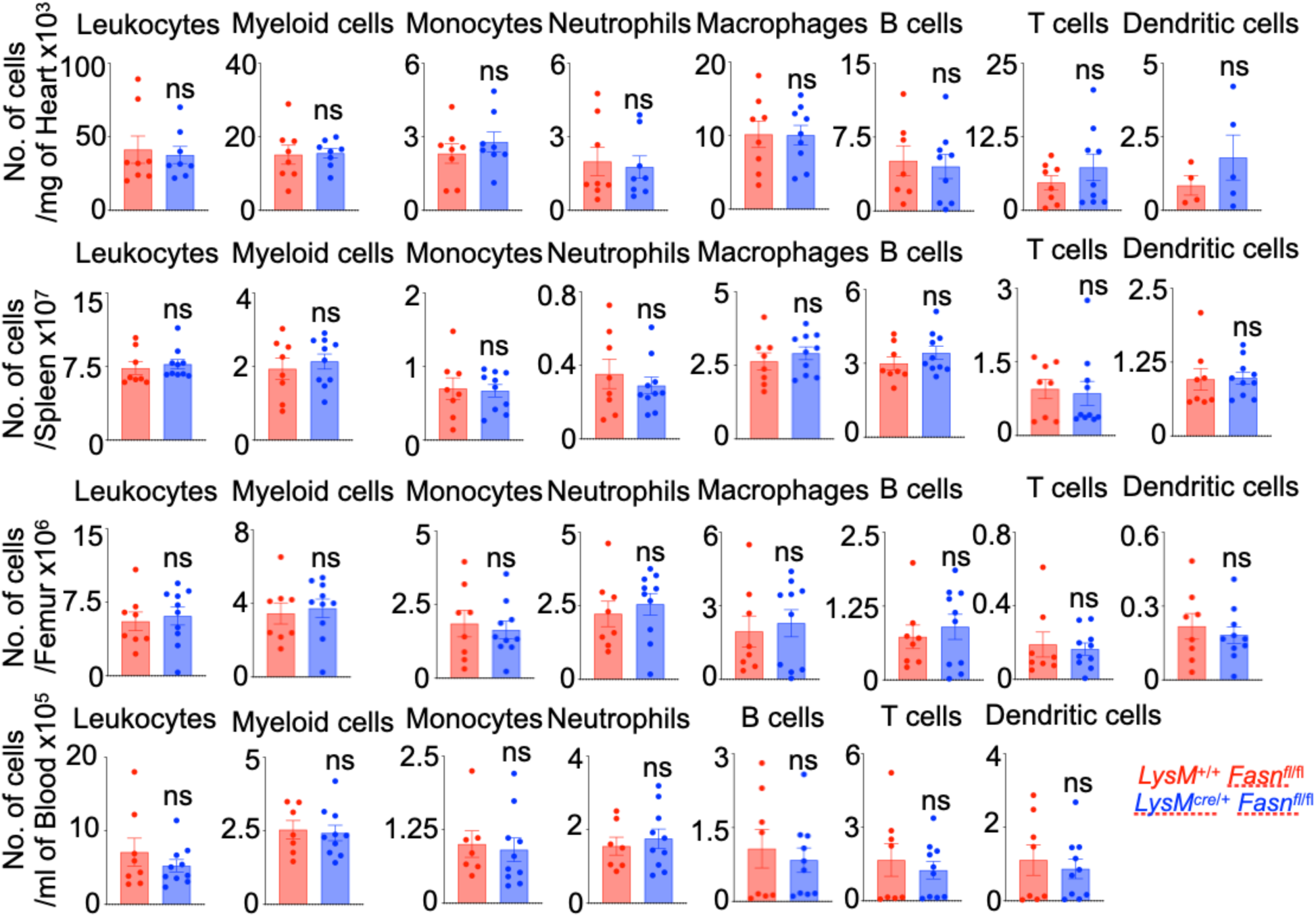
Myeloid *Fasn* deficiency does not alter leukocyte subset numbers after MI. Quantification of different leukocyte subsets in the heart, spleen, bone marrow, and blood in *LysM*^+/+^ *Fasn*^fl/fl^ and *LysM^cre/+^ Fasn*^fl/fl^ 28 days after MI were caried out by flow cytometry (n=7-10/group). The data are expressed as mean ± SEM and pooled from 3–5 independent experiments. The Mann-Whitney U test was performed to determine statistical significance.

**Supplemental Figure 4:**
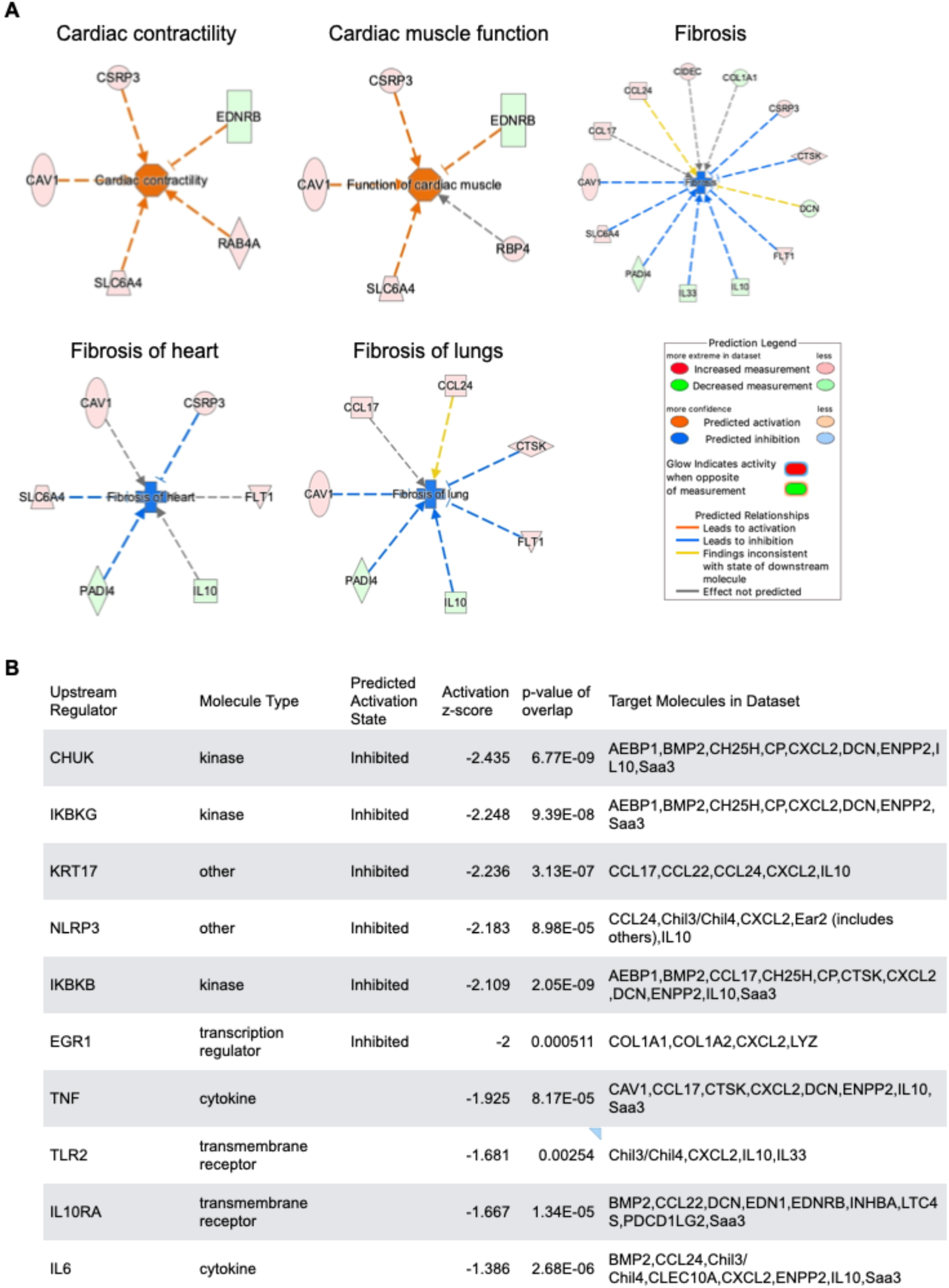
*Acly* deficiency in cardiac macrophages reduces the enrichment of the genes involved in organ fibrosis. Cardiac macrophages were isolated from *LysM*^+/+^ *Acly*^fl/fl^ and *LysM^cre/+^ Acly*^fl/fl^ mice on day 28 after MI, and bulk RNA sequencing was performed. **A.** The network diagrams show the top five pathways altered in *LysM^cre/+^ Acly*^fl/fl^ cardiac macrophages compared to *LysM*^+/+^ *Acly*^fl/fl^ cardiac macrophages (n=3). **B.** The table lists the top 10 upstream regulators *LysM^cre/+^ Acly*^fl/fl^ cardiac macrophages (n=3). The data from an RNA sequencing experiment are shown.

**Supplemental Figure 5:**
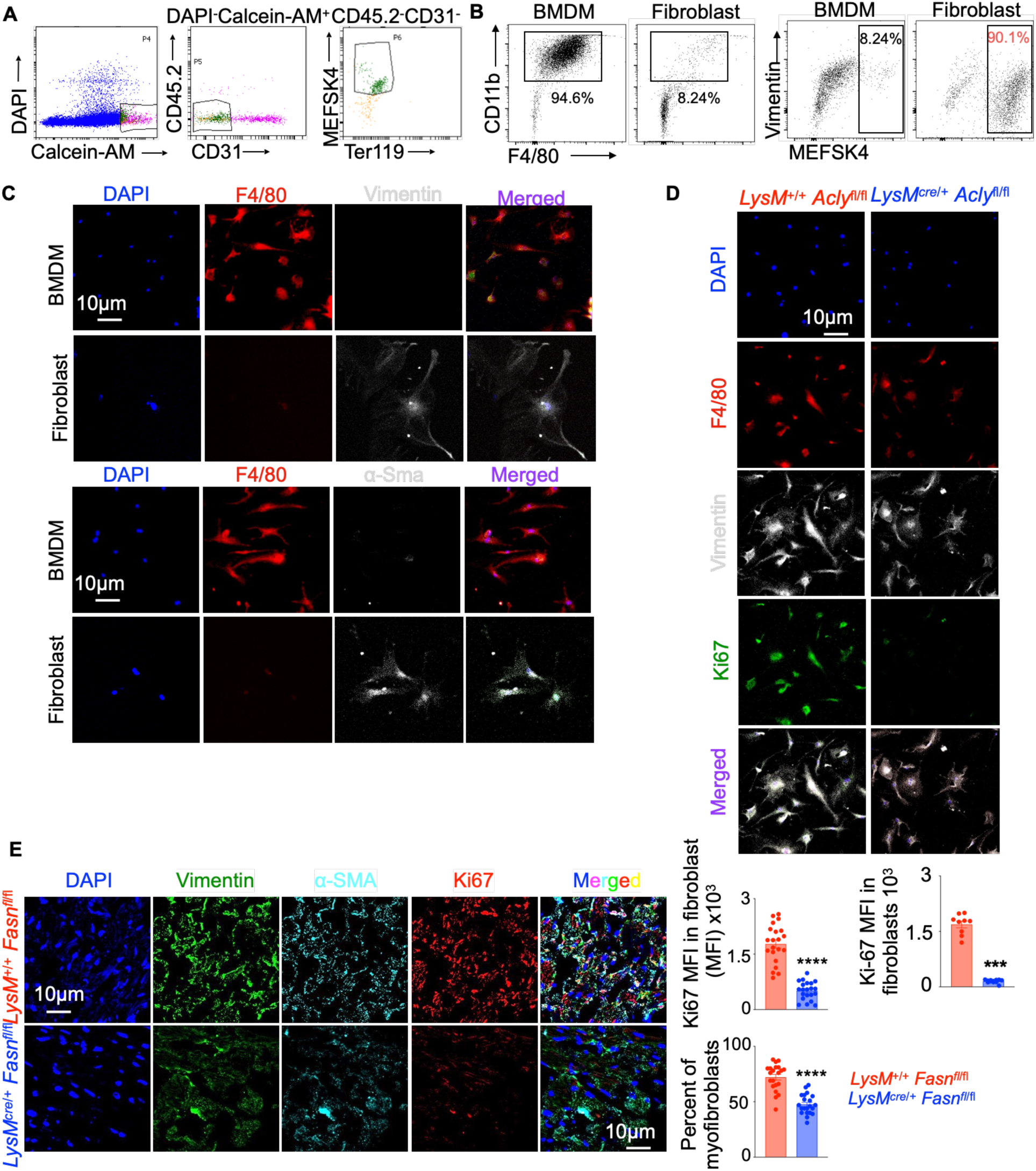
Macrophage ACLY expands myofibroblasts *in vitro*. A. Flow cytometric gating strategies to isolate cardiac fibroblasts from the hearts of *LysM*^+/+^ *Acly*^fl/fl^ and *LysM^cre/+^ Acly*^fl/fl^ mice on day 28 after MI are provided. **B-C.** The representative flow cytometry plots (**B**) and immunofluorescence images (**C**) show the purity of BMDM (CD11b^+^ F4/80^+^, left) and cardiac fibroblasts (vimentin^+^ MEFSK4^+^, right). (n=5-7/group) **D-E.** C57BL/6 cardiac fibroblasts were cocultured with BMDM isolated from *LysM*^+/+^ *Acly*^fl/fl^ and *LysM^cre/+^ Acly*^fl/fl^ mice. Fibroblast proliferation was analyzed by confocal microscopy (n=5-7/group). The data are expressed as mean ± SEM and pooled from 3–5 independent experiments. The Mann-Whitney U test was performed\*\*\**p* < 0.001.

**Supplemental Figure 6:**
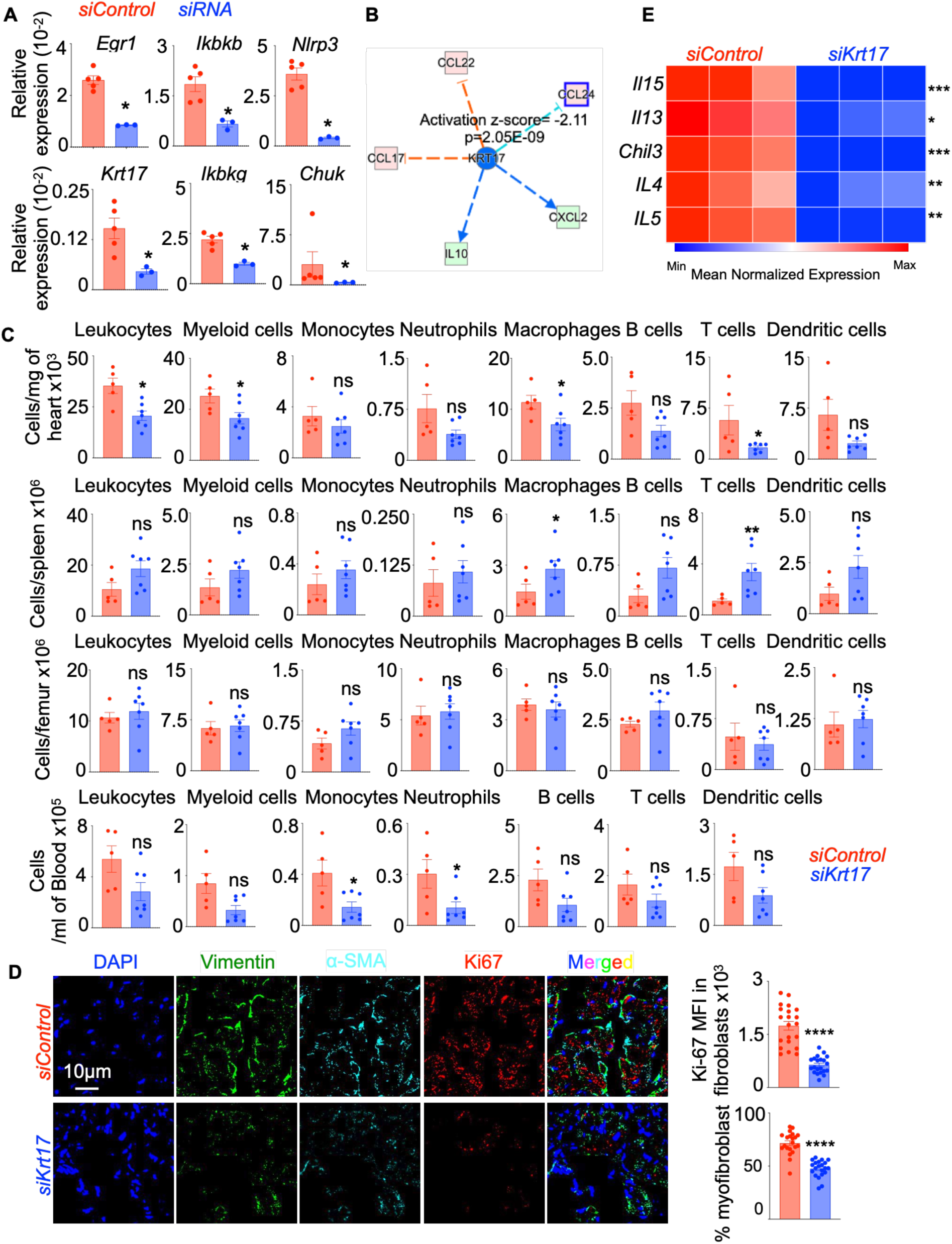
Myeloid Krt17 leads to augmented fibroblast proliferation in the heart after MI. **A.** qPCR was performed to validate the silencing of *Egr1*, *Ikbkb*, *Nlrp3*, *Krt17*, *Ikbkg*, and *Chuk* in BMDM (n=6-10/group). **B.** The network diagram lists the top 5 targets of Krt17 using an IPA analysis of the differentially expressed genes in *LysM^cre/+^ Acly*^fl/fl^ cardiac macrophages compared to *LysM*^+/+^ *Acly^fl/fl^* cardiac macrophages (n=3). **C-D.** C57BL/6 mice were treated with si*Control* and si*Krt17* incorporated in lipidoid nanoparticles, which are preferentially engulfed by macrophages, for 28 days after MI (n=5-7/group). The immune cells enumerated by flow cytometry in the heart, spleen, bone marrow, and blood (**C**). Confocal microscopy was employed to quantify fibroblast proliferation and myofibroblasts frequency in the heart (**D**). **E.** The heatmap demonstrates the expression of the pro-fibrotic genes in *siControl* and *siKrt17*-treated BMDM (n=5-7/group). The data are expressed as mean ± SEM and pooled from 3–5 independent experiments. The Mann-Whitney U test was performed. \**p* < 0.05, \*\**p* < 0.01, \*\*\**p* < 0.001, \*\*\*\**p* < 0.0001.

**Supplemental Figure 7:**
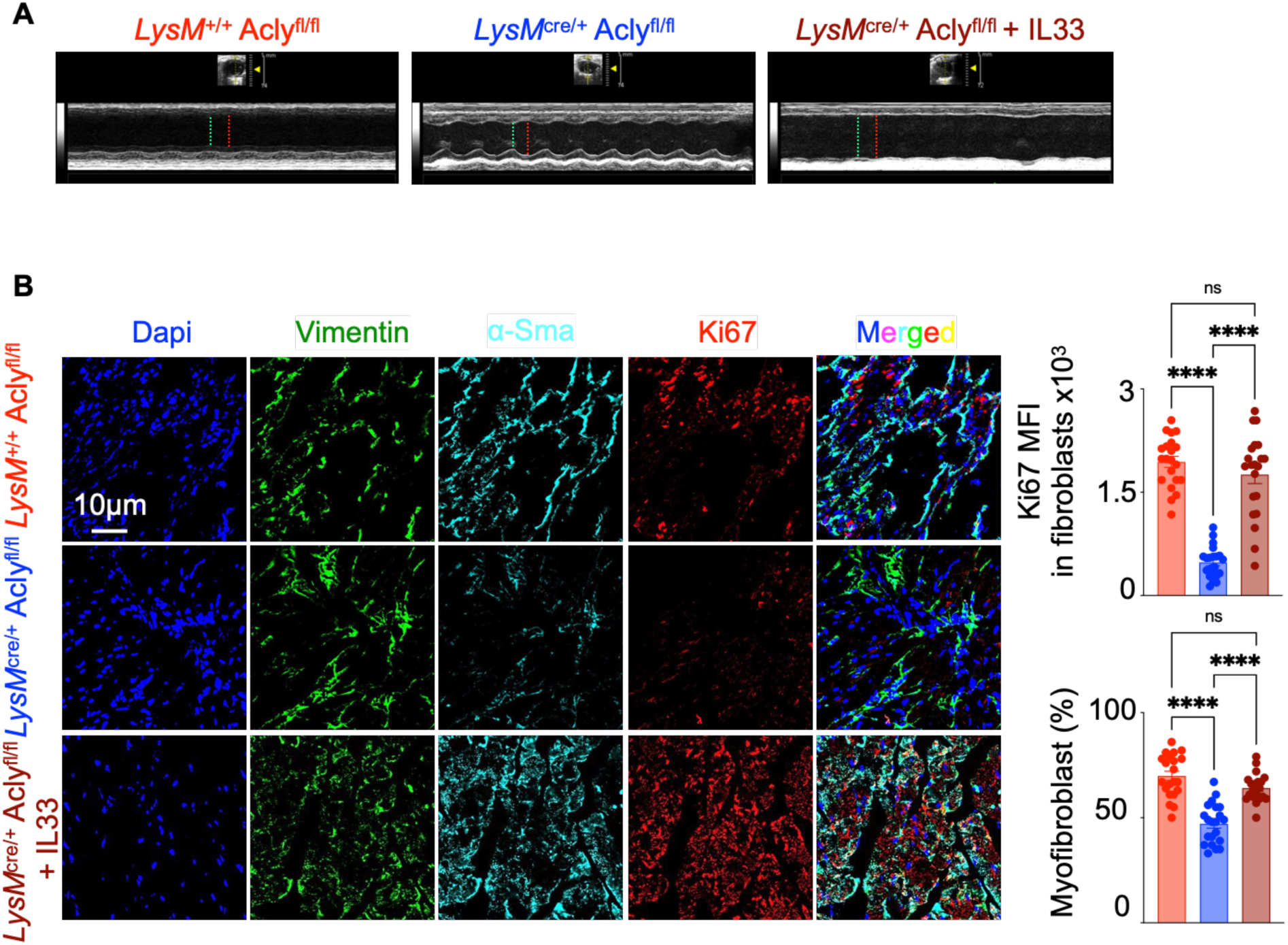
IL-33 supplementation restores cardiac fibrosis in mice lacking myeloid *Acly*. **A-B.** *LysM*^+/+^ *Acly*^fl/fl^ and *LysM^cre/+^ Acly*^fl/fl^ mice were treated with either vehicle or rIL-33 for 28 days post MI (n=9-6/group). The representative echocardiograms in M-mode of the LV in SAX view obtained by echocardiography are included. The green and red lines indicate LVID;s = left ventricular internal diameter in systole and LVID;d = left ventricular internal diameter in diastole respectively(**A**). Immunofluorescence images show vimentin, α-SMA, and Ki67 staining, and the bar graphs display cardiac fibroblast proliferation (Ki-67^+^) and myofibroblasts proportion (**B**). The data are expressed as mean ± SEM and pooled from 3–5 independent experiments. \*\*\*\**p* < 0.0001. One way ANOVA was performed.

**Supplemental Figure 8:**
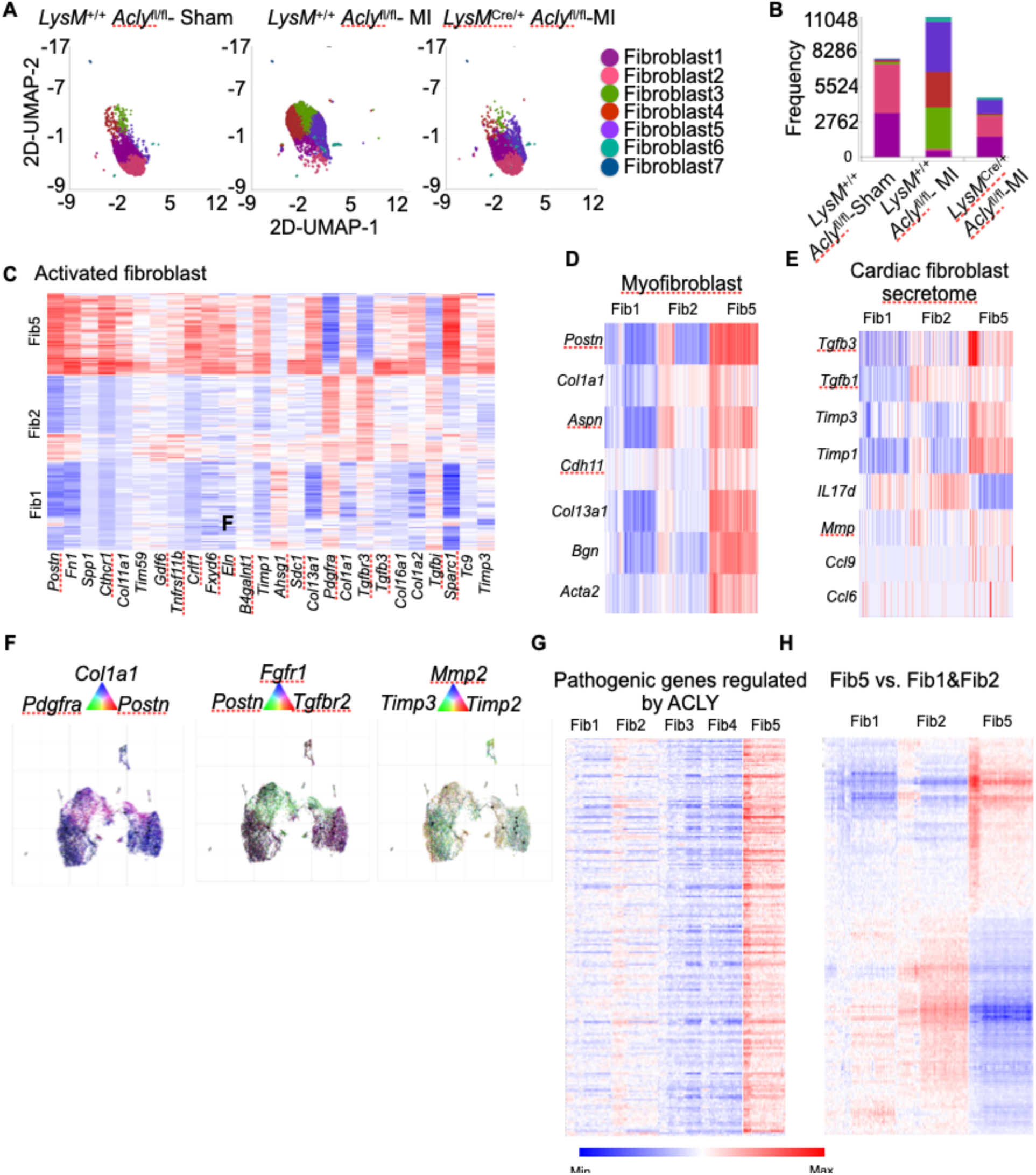
Macrophage ACLY promotes Fibroblast 5 expansion after MI. Cardiac fibroblasts were isolated from the heart of *LysM*^+/+^ *Acly*^fl/fl^ mice with sham and MI surgeries and *LysM*^Cre/+^ *Acly*^fl/fl^ mice with MI on day 28 after the surgery. Single cell RNA sequencing was performed on the sorted cells. n=4/ group. **A-B.** The UMAP plots (**A**) and bar graphs (**B**) demonstrate the distribution of different fibroblasts subtypes. **C-E.** The heatmaps depicts the expression of the genes specific for activated fibroblasts (**C**), myofibroblasts (**D**), and cardiac fibroblast secretome (**E**) in Fibroblasts 1, 2, and 5. **F.** The pseudotime trajectory plots in the numeric triads displays the expression of the genes specific to activated fibroblasts and their secreting factors (*Col1a1*, *Pdgfra*, *Postn*, *Fgfr1*, *Tgfbr2*, *Mmp2*, *Timp2,* and *Timp3*) **G-H.** The heatmaps reveals the expression of the genes regulated by ACLY (**G**) and differentially expressed genes in Fibroblast 5 compared to Fibroblasts 1&2 (**H**) in the various fibroblast subsets.

## Notes

**Declaration of interest:** The authors have declared that no conflict of interest exists.

### Competing Interest Statement

The authors have declared no competing interest.

